# Analyzing the intrastate and interstate swine movement network in the United States

**DOI:** 10.1101/2024.01.25.576551

**Authors:** Nicolas C. Cardenas, Arthur Valencio, Felipe Sanchez, Kathleen C. O’Hara, Gustavo Machado

**Affiliations:** Department of Population Health and Pathobiology, College of Veterinary Medicine, North Carolina State University, Raleigh, North Carolina, USA; Center for Geospatial Analytics, North Carolina State University, Raleigh, NC, USA; U.S. Department of Agriculture, Animal and Plant Health Inspection Service, Veterinary Services, Strategy and Policy, Center for Epidemiology and Animal Health, Fort Collins, CO, USA

**Keywords:** Pig movement, multistate animal transportation, contact chain, swine network

## Abstract

Disease prevention and control tactics rely on identifying and restricting animal movement to attenuate the between-premises spread of disease in livestock systems. Therefore, it is essential to uncover between-premises movement dynamics, including shipment distances and network-based control strategies. Here, we analyzed three years of between-premises pig movements, which include 197,022 unique animal shipments, 3,973 premises, and 391,625,374 pigs shipped across 20 U.S. states. We constructed unweighted, directed, temporal networks at 180-day intervals to calculate premises-to-premises movement distances, the size of connected components, network loyalty, and degree distributions, and, based on the out-going contact chains, identified network-based control actions. Our results show that the median distance between premises pig movements was 74.37 km, with median intrastate and interstate movements of 52.71 km and 328.76 km, respectively. On average, 2,842 premises were connected via 6,705 edges, resulting in a weak giant connected component that included 91% of the premises. The premises-level network exhibited loyalty, with a median of 0.65 (IQR: 0.45 – 0.77). Results highlight the effectiveness of node targeting and disease spread; we demonstrated that targeting 25% of farms with the highest degree or betweenness limited spread to 1.23% and 1.7% of premises, respectively. While there is no complete shipment data for the entire U.S., our multi-state movement analysis demonstrated the value and the needs of such data for enhancing the design and implementation of proactive-disease control tactics.

## 1. Introduction

The shipment of animals between premises has been widely recognized as the primary route involved in the transmission of infectious pathogens of swine such as African swine fever (ASF), porcine respiratory and reproductive syndrome virus (PRRSV), and foot-and-mouth disease (FMD) (Galvis et al., 2022a; Passafaro et al., 2020; Vinueza et al., 2022; Hammami et al., 2022a; Cardenas et al., 2022, 2021; Galvis et al., 2021; VanderWaal et al., 2020; Lee et al., 2017; White et al., 2017; Neumann et al., 2005; Pappaioanou and Gramer, 2010; Fèvre et al., 2006; Sykes et al., 2023). Disease intervention strategies involve reducing the number of between-premises animal movements; therefore, elucidating local and nationwide movement network patterns and identifying shipment hubs and road pathways used for pig shipment are expected to provide essential information about disease spread potential and efficient outbreak control (Puspitarani et al., 2023; Passafaro et al., 2020; Cardenas et al., 2022; Hammami et al., 2022a; Gorsich et al., 2019; Strano et al., 2018; Sykes et al., 2023.)

The U.S. swine industry is vertically integrated and comprised of approximately 64,871 premises, housing 72 million pigs in premises specializing in one or more production phases (e.g., farrow-to-wean, growing pig) (Passafaro et al., 2020; NASS et al., 2022; USDA and NASS, 2019; Reimer, 2006). In North America, the movement of pigs occurs in groups (all-in / all-out) or as needed (continuous flow) (Passafaro et al., 2020; NASS et al., 2022; USDA and NASS, 2019; Reimer, 2006). An estimated 71% of growing pigs in the U.S. are finished in a different location than they were farrowed (Cabezas et al., 2021). Moreover, animals moved from breeding farms to different premises (e.g., nurseries, wean-to-feeder) are mixed into populations that often contain animals originating from multiple premises, known as commingling (Nadal-Roig et al., 2020). Commingling has been previously linked to a heightened risk of disease transmission (Aragon et al., 2019). Movement flows that rely on many premises as sources to fill nursery or finisher premises result in contact networks with low animal movement loyalty, leading to a reduction in the number of regular connections between the same premises, thus increasing the chances for disease dissemination (Puspitarani et al., 2023; Hammami et al., 2022b; Schulz et al., 2017.)

Previous studies have described U.S. swine contact networks by examining network loyalty and movement distances by measuring geodesic distance (Kinsley et al., 2019; Makau et al., 2021; Moon et al., 2019) and between-premises movement patterns by characterizing the network structure, analyzing connected components, and identifying potential targets for disease control (Passafaro et al., 2020; Cabezas et al., 2021; Passafaro et al., 2020; Lee et al., 2017). However, analyses using multi-state swine movement network data that examine intrastate and interstate movement patterns have been limited due to the lack of a centralized electronic system that captures and monitors swine movement records (Gorsich et al., 2019). More importantly, the lack of movement data recording standards and centralized movement data repositories has led to attempts to predict shipment patterns within states (Moon et al., 2019) and nationally (Passafaro et al., 2020) by scaling up available movement data.

The extraordinary complexity of between-premises animal movements, which change daily (Galvis et al., 2022a; Sykes et al., 2023), presents a formidable challenge for the swine industry and decision-makers who must implement effective disease control measures under time constraints. A European study that used real pig movement data and network modeling has demonstrated that between-farm movements may account for a staggering 99% of ASF transmission among commercial herds (Andraud et al., 2019). A more recent U.S. model simulation study showed that between-farm movement of swine was the predominant route of transmission of ASF, with an average contribution of 71.4% (Sykes et al., 2023). Given these results, an in-depth examination of real swine contact networks across multiple states would provide invaluable insights for developing response and recovery tactics.

To address the lack of comprehensive movement data and to better prepare for disease outbreaks, we formed a consortium of multiple states and sectors to develop epidemiological models to combat swine infectious diseases and other animal health threats (Machado et al., 2023). Leveraging the consortium, we have amassed multi-state animal movement data and present a large-scale network analysis of between-premises shipments from 3,973 premises, moving 391,625,374 pigs across 20 U.S. states and six production companies. Our analysis compares the number of animal movements by premises’ production types and swine production companies, investigating premise-to-premise intrastate and interstate shipment distances and the mixing of movements at the destination premises. Finally, we used a temporal network to examine the effect of node removal in curbing disease spread trajectories.

## 2. Material and methods

### 2.1 Swine movement data

A total of 282,185 premises-to-premises records were obtained from six U.S. swine production companies representing movements occurring between January 1, 2020, and January 15, 2023. The movement data included unique premises identification numbers, geolocations, premises of origin and destination, production type (i.e., finisher, nursery, sow), movement date, and the number of pigs transported. Of these records, 57,237 were discarded due to missing data associated with premises identification or movement date. Additionally, 60,022 movements to slaughterhouses were discarded, as we consider those movements as epidemiological endpoints. The final dataset included 197,022 unique animal shipments among 3,973 premises and 391,625,374 pigs shipped across 20 U.S. states: Missouri, Iowa, Wyoming, Illinois, Nebraska, Oklahoma, Indiana, North Carolina, Mississippi, Kansas, Texas, Colorado, Virginia, South Carolina, Pennsylvania, Ohio, New York, Arizona, Utah, and Michigan. The distribution of pig movements by the state is shown in Figure 1 and by the production company in Supplementary Material, Figure S1.

**Figure 1.**
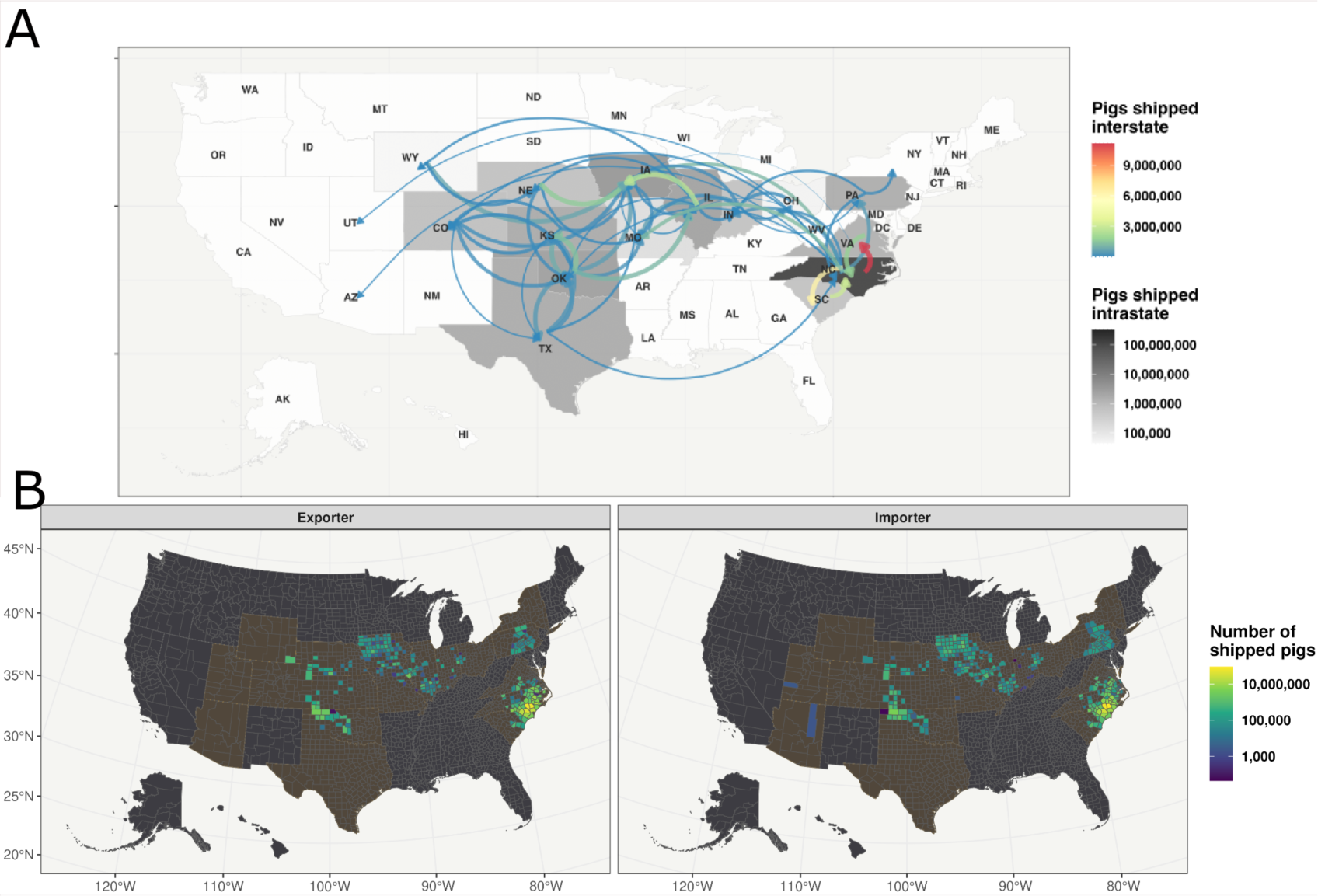
The intrastate and interstate and between counties movement of pigs. A) The shaded gray background shows the number of intrastate pig shipments, while the color and width of the arrow lines represent the interstate shipments. B) Exporter counties represent out-going pig movements, while the importer panel represents incoming pig movements from January 1, 2020, to January 15, 2023. The number of shipped pigs is on a log10 scale, and states with at least one shipment are highlighted.

### 2.2 Premises information

Premises that received or sent at least one animal during the timeframe of this study were grouped into six categories based on swine production types along with movement records. These categories were: Sow premises: Premises with breeding age, gestation units, or farrowing rooms. Nursery premises: Premises that raise piglets from weaning (approximately three weeks of age) to about ten weeks of age. Finisher premises: Premises that raise pigs from approximately ten weeks of age until reaching market weight at five to six months. Wean-to-finish premises (wean-to-finisher): Premises that raise pigs from weaning age to market weight. Gilt developer unit (GDU): Breeding premises specialized in gilt development. Boar stud: Premises with male pigs of reproductive age. The criteria for grouping premises by type are presented in Supplementary Material Table S2. Of the 2,595,404 premises, 10.16% reported more than one production type. The total number of premises with at least one animal shipment during the study period is shown in Table 1.

**Table 1.** The number of active premises in each company, grouped by production type.

| **Company** | **Finisher** | **Nursery** | **Sow** | **Wean-to-finisher** | **Boar stud** | **Total** |
| --- | --- | --- | --- | --- | --- | --- |
| A | 1,117 | 389 | 234 | 257 | 0 | **1,997** |
| B | 198 | 53 | 58 | 2 | 1 | **312** |
| C | 89 | 39 | 24 | 42 | 0 | **194** |
| D | 649 | 180 | 87 | 0 | 6 | **922** |
| E | 210 | 59 | 31 | 0 | 0 | **300** |
| F | 332 | 40 | 64 | 183 | 0 | **619** |
| **Total** | **2,595** | **760** | **498** | **484** | **7** | **4,344** |

### 2.3 Intrastate and interstate movements

We used the premises geolocations to up-scale between-premises movements and calculate the number of movements within and between counties and states. For state-level movements, we calculated the number of intrastate and interstate shipments and the number of shipped pigs. The results are depicted in thematic maps (Figure 1). Counties were also mapped based on their movement pattern profile; counties that were primary sources of pigs were labeled “exporters,” while counties that were primary destinations of pigs were labeled “importers.”

#### 2.3.1 Between-premises shipment analysis

The shortest road paths between premises were calculated using the premises’ geolocations and the OpenStreetMap® calculation paths service (Giraud, 2022; Luxen and Vetter, 2011). The distance of 25,524 unique pairs of origin-destination shipments was calculated. For all computed pathways, the average number of shipments and animals for each road segment was calculated. In addition, the distance from the source to the destination premises was calculated, considering the distances of intrastate and interstate movements for each company and production type. Finally, the distances were transformed into an empirical cumulative distribution function (eCDF) (Ayer et al., 1955). An eCDF plot shows the data from the lowest to the highest values and compares them to their percentiles, allowing us to display the cumulative distribution probability observed over the estimated distances (Ayer et al., 1955.)

### 2.4 Mixing of animals at destination premises

Here, we use commingling to refer to any mixing of animals at the destination premises, regardless of production phase (e.g., at nurseries, finishers, and sow farms). To quantify the amount of mixing sources, we used a premises-level network that considered the origin and destination production type. A contact network was constructed by looking at the ingoing and outgoing movements of each premises over 180 days windows (described in section 2.5). A one-way Kruskal-Wallis test was conducted to compare commingling among production types and companies, followed by Dunn’s multiple comparison post hoc test (Kruskal and Wallis, 1952.)

### 2.5 Dynamic network analysis

Unweighted directed temporal networks were constructed with nodes representing each premises and edges representing animal movements between premises. Each time frame of the temporal network is represented by movements occurring over a 180-day interval, which is an astable representation of pig movement network parameters (Makau et al., 2021). The time step between each of the frames of the temporal network was defined as seven days, which coincides with the frequency of most of the animal movements’ information provided by participating companies. This means the first time frame we analyzed was at *t_0_*= 01/January/2020, with a time window of Δ*t_1_’*=01/January/2020 to 29/June/2020. The next time window was from Δt1’ = 08/January/2020 to 06/July/2020. The number of nodes, edges, network diameter, cluster coefficient, associativity, giant weak connected component (GWCC), and giant strongly connected component (GSCC) was computed for each company, generating a time series of these parameter values and summarizing the mean and the 95% confidence intervals. The definitions for each of the network parameters are provided in Supplementary Material Table S1.

#### 2.5.1 Dynamic premises-level network degree distribution

The scale-free hypothesis is a concept from network theory that describes a specific property of complex networks in which a few nodes (premises) have a significantly higher number of connections (degree) than most nodes in the network. This phenomenon is particularly relevant in disease transmission, as these highly connected nodes can serve as major sources of contagion. Here, animal movements between premises were split into 180-day time-window intervals for the entire study period for each company, and production types were used to fit in-degree and out-degree distributions from each time frame. The scale-free hypothesis was tested by fitting the distribution of degrees *k* to a power law, defined as *p*(*k*) = *ak*^−α^, where α is the scale parameter and *a,* a proportionality term (Barabási et al., 1999; Passafaro et al., 2020). The data points used in these analyses were averages and standard deviations of each degree value across premises. In-degree and out-degree distributions were fitted to a range of distributions to compare the fit to the data and theoretical power-law distributions. We used goodness-of-fit tests based on the Kolmogorov-Smirnov statistic and calculated log-likelihood ratios between the candidate distributions. We used p-values to determine the significance of the results over 10,000 bootstrapping simulations. In-degree and out-degree distributions fit a power law if the p-value exceeds 0.05 (Clauset et al., 2009.)

#### 2.5.2 Network loyalty

Loyalty was defined as the fraction of edges preserved in two consecutive time-frames, calculated by the Jaccard index, which provides a value from 0 to 1. A loyalty of 0 denotes all edges having changed in the consecutive time frame(s), and a loyalty of 1 represents all edges remaining the same (Schulz et al., 2017; Valdano et al., 2015). Here, we examine between-premises directed temporal network metrics, including incoming and outgoing edges by production type.

Briefly, production-type loyalty for type *i* expresses the fraction of edges preserved between consecutive time frames for premises of type *i* and premises to which they have moved into or from. For example, sow in-loyalty can take a value between 0 and 1, expressing how many movements from premises {1, 2, 3, …} to sow premises remain the same in two consecutive time frames of the temporal network. Mathematically, the production-level loyalty is defined as

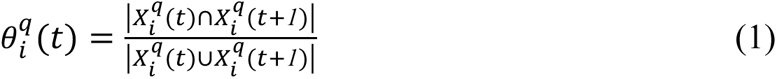

where *t* is the current time window (180 days), *t*+1 corresponds to the consecutive time window (180 days), *q* = {*in*, *out*} and is an index defining if the link being analyzed is of production type *i* sending animals to the set *X* (i.e., *q* = *out*) or production type *i* receiving animals from the set *X* (i.e., *q* = *in*). *X* is the set of premises linked to any premises of production type *i* (i.e., set of premises {1, 2, 3, …} that have movements into (in-loyalty) or from (out-loyalty) production type *i*). The production-type loyalty distributions and time series have been calculated for the overall movement data and each company (Supplementary Material Figure S12.)

### 2.6 Between-premises cumulative contacts as a proxy of outbreak size and effectiveness of network-based targeted percolation

Network percolation is a method that approximates the effects of countermeasure actions, such as farm closure or movement restrictions, that effectively reduce disease dissemination by targeting premises based on their network role (Dubé et al., 2008; Nöremark et al., 2011; Rossi et al., 2017). To identify the most effective network-based target interventions, we calculated four network parameters for each premise: degree, betweenness, cluster coefficient, and PageRank (Supplementary Material Table S1). Once the premises were ranked based on each network parameter, percolation was used to remove nodes (i.e., premises) one at a time, to include up to 5%, 10%, 15%, 20%, and 25% of the total number of premises. The percentage removal was proportional to the number of premises in each production company. For example, for company A, 5% removal was 100 out of 1,997 premises. Percolation was also performed without any premises ranking. This scenario was used to mimic regular random surveillance activities and to benchmark with network-based interventions. To examine the expected reduction of outbreak size achieved by targeting premises based on their network roles, we adapted the method proposed by (Payen et al., 2019) to calculate the out-going contact chains as a proxy for expected outbreak size, named hereafter as “spread cascade model” (Cardenas et al., 2021; Machado et al., 2021; Payen et al., 2019). Here, spread cascade size reduction was calculated for the percolation scenario; company-level results are also available in the supplementary material.

### 2.7 Software

This study was conducted in R statistical software v. 4.1.1 (R Core Team, 2021) using the following packages: tidyverse (Wickham et al., 2019), sf (Pebesma, 2018), EpiContactTrace (Nöremark and Widgren, 2014), igraph (Csardi and Nepusz, 2006), osrm (Giraud, 2022), poweRlaw (S. Gillespie, 2015), and lubridate (Grolemund and Wickham, 2011.)

## 3. Results

### 3.1 Intrastate and interstate movements

The daily number of shipped pigs exhibited a median of 356,848 (interquartile range [IQR]: 57,350-468,642, maximum: 2,359,236) and 156 shipments (IQR: 41-311, maximum: 453). The states with the highest intrastate movements were North Carolina, Iowa, and Oklahoma, with 88,749, 17,505, and 15,717 shipments, and 322,713,497; 7,001,558, and 4,861,502 shipped pigs, respectively (Figure 1). North Carolina, Iowa, and Oklahoma also had the highest number of between-counties pig movements, with a median of 6,434 (IQR: 2,386-23,100, maximum: 20,569,392) and 30,653 (IQR: 5,704-202,464, maximum: 51,099,957), respectively. The top three states with the highest between-state movements were North Carolina to Virginia, with 3,011 shipments and 11,152,229 pigs; North Carolina to South Carolina, with 2,282 shipments and 6,311,509 pigs; and Illinois to Iowa, with 1,510 shipments and 3,038,413 pigs (Figure 1).

Results of interstate and intrastate movements by production types are shown in Supplementary Material Figures S3-S7.

### 3.2 Between-premises shipment analysis

Road pathways linking North Carolina, Virginia, and Iowa contained the highest quantity of between-premises movements (Figure 2 and Supplementary Material Figure S8.)

**Figure 2.**
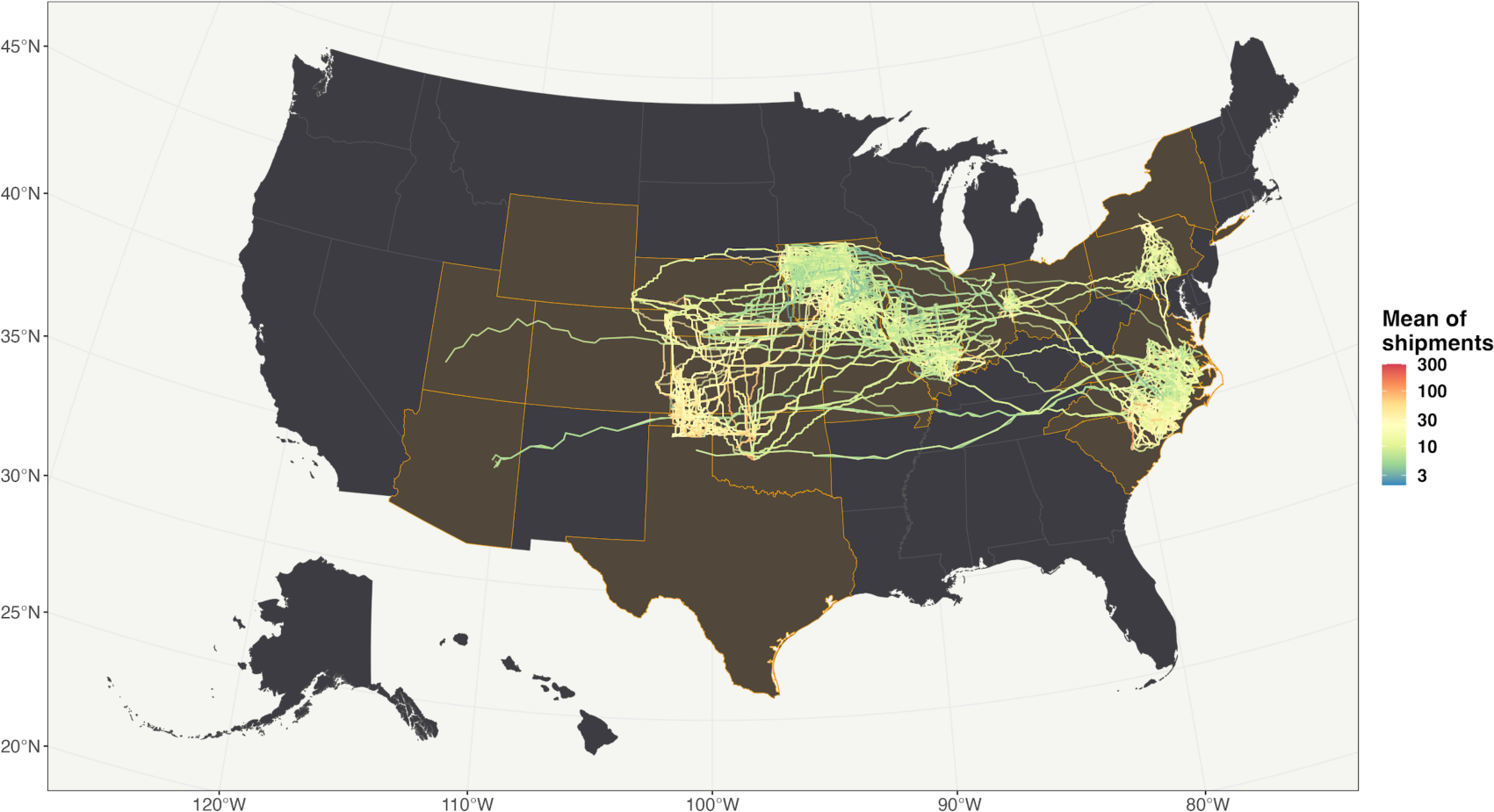
Between premises shipment. The lines represent the between-premises pathways. The color represents the mean number of shipments from January 1, 2020, to January 15, 2023. States with at least one shipment are highlighted.

The median distance pigs were shipped was 74.37 km (IQR: 33.79-187.76 km, maximum: 3,415.22 km) (Figure 3A). Intrastate movements had a median distance of 52.71 km (IQR: 26.24-104.15 km, maximum: 534.00 km), while interstate movements had a median distance of 328.76 km (IQR: 191,438-705,680 km, maximum: 3,415.22 km) (Figure 3B). Among production types, boar stud shipments were by far the most distant, with a median of 237 km (IQR: 148-1,026 km, maximum: 1,162 km). Wean-to-finisher shipments had a median distance of 141 km (IQR: 51.5-285 km, maximum: 1,810 km), and finisher shipments had a median distance of 124 km (IQR: 51-231 km, maximum: 3,099 km) (Figure 3C.)

**Figure 3.**
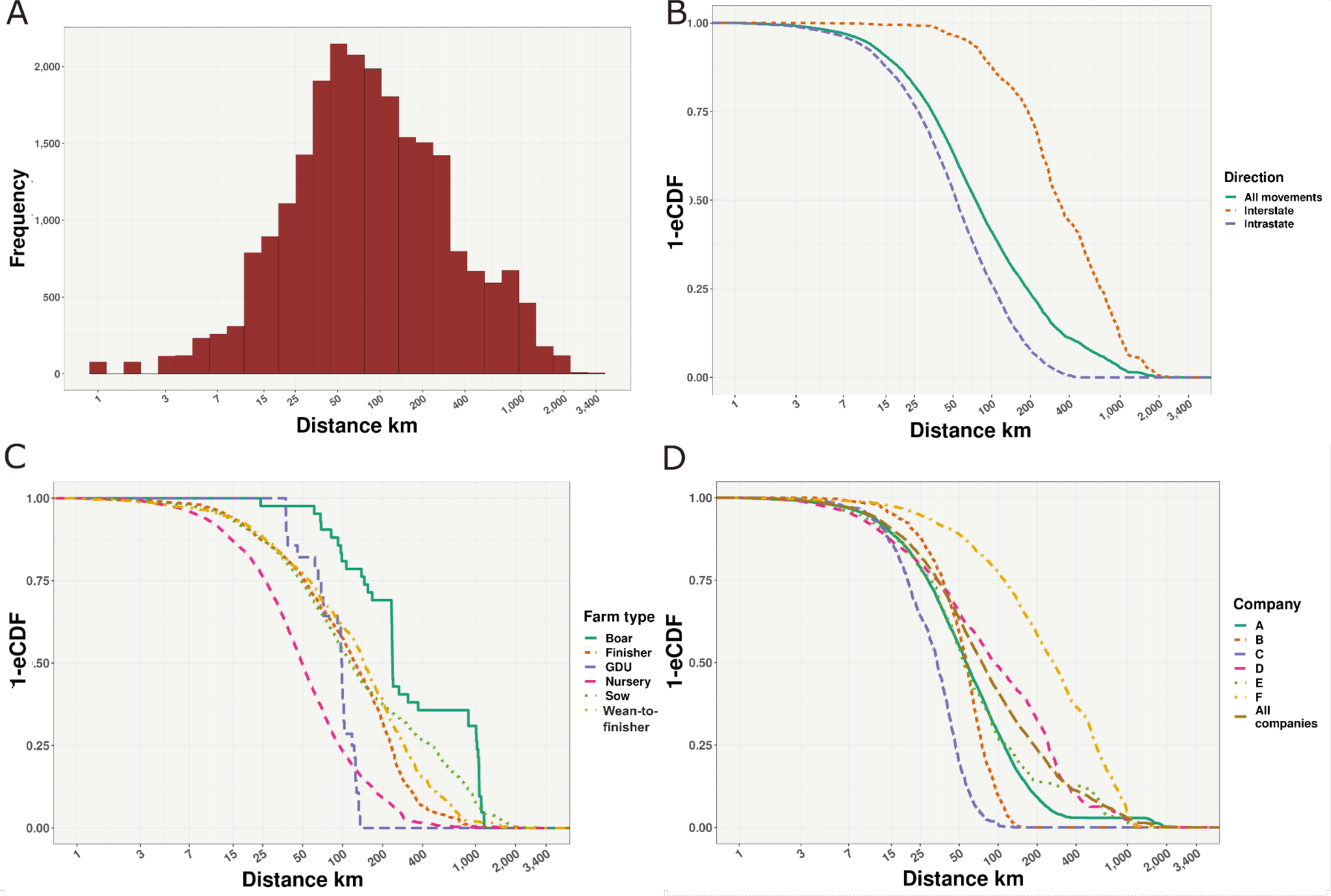
Between-premises movement distances. A) The histogram of between-premises shipment distances. B) Intrastate and interstate empirical cumulative distribution function (eCDF). C) The eCDF by production type. D) The eCDF by company.

Comparing shipment distances, companies (A, E, and C) shipments were at short distances, with median distances of 56.1 km (IQR: 28.4-11 km, maximum: 3,415 km), 54.1 km (IQR: 30.5-113 km, maximum: 1,046 km), and 33.7 km (IQR: 19.8-46.6 km, maximum: 132 km), respectively. However, companies F, D, and B shipments were long distances, with median distances of 264 km (IQR: 108-567 km, maximum: 1,603 km), 91.7 km (IQR: 33.6-245 km, maximum: 1,250 km), and 56.8 km (IQR: 37.4-74.8 km, maximum: 166 km), respectively (Figure 3D). Distributions of road distances for each company by production types are shown in Supplementary Material Figures S9 and S10.

### 3.3 Number of mixing animals at destination premises (commingling)

Regardless of production type, the median number of different premises sources mixing at destination premises was 2 (IQR: 1-4, maximum: 144). Commingling in wean-to-finisher premises was higher than in finisher, sow, and nursery farms, respectively (Table 2). Different premises sending pigs from sow into wean-to-finisher exhibited a median of 4 (IQR: 2-7, maximum: 24), and sow sending pigs into finisher had a median of 4 (IQR: 2-10, maximum: 22), followed by sow to finisher and finisher to sow farms with a median of 3 (IQR: 2-6, maximum: 24) and median of 3 (IQR: 2-5, maximum: 21), respectively. The largest commingling values were observed for movements between finisher premises and finisher to nursery premises, with up to 144 different premises mixing pigs at destination premises (Table 2 and Supplementary Material Table S4 and Figure S11). Finally, we compared the levels of commingling by farm type among companies and noted that 73.33% of the pair comparisons were significantly different (Supplementary Material Table S4 and Figure S12.)

**Table 2.** The median and interquartile range of pig mixing (commingling) - number of different sources of pigs entering the premises within a given time 180 days window - by origin production type from 1st January 2020 to 15th January 2023.

| Origin | Destination | Number of movements | Median | 25% quantile | 75% quantile | Maximum |
| --- | --- | --- | --- | --- | --- | --- |
| Finisher | Finisher | 733 | 2 | 1 | 6 | 144 |
| Finisher | Nursery | 328 | 1 | 1 | 2 | 144 |
| Sow | Finisher | 487 | 3 | 2 | 6 | 24 |
| Sow | Wean-to-finisher | 507 | 4 | 2 | 7 | 24 |
| Nursery | Finisher | 2059 | 2 | 2 | 4 | 24 |
| Nursery | Wean-to-finisher | 78 | 1.5 | 1 | 3 | 24 |
| Finisher | Wean-to-finisher | 325 | 4 | 2 | 10 | 22 |
| Finisher | Sow | 497 | 3 | 2 | 5 | 21 |
| Sow | Nursery | 738 | 2 | 1 | 4 | 20 |
| Nursery | Nursery | 56 | 2 | 1 | 6 | 12 |
| Wean-to-finisher | Finisher | 237 | 1 | 1 | 2 | 6 |
| Wean-to-finisher | Nursery | 28 | 1 | 1 | 2.5 | 6 |
| Wean-to-finisher | Sow | 45 | 2 | 1 | 2 | 6 |
| Wean-to-finisher | Wean-to-finisher | 129 | 2 | 1 | 2 | 6 |
| Nursery | Sow | 47 | 2 | 1 | 3 | 6 |
| Sow | Sow | 161 | 2 | 1 | 2 | 4 |

### 3.4 Dynamic network analysis

The weekly network of pig movements shown an average of 2,842 premises receiving and or sending pigs and an average of 6,705 edges (Table 3). The average number of premises per company receiving or sending at least one shipment per week ranged from 171.25 to 1,586. The network diameters across companies were not significantly different, varying from 7.7 to 11.84. However, network assortativity was significantly different among companies. Companies F, A, and E exhibited positive assortativity from 0.56 to 0.03, while D, C, and B exhibited negative values from -0.24 to -0.52. Overall, GSCC ranged from 0.2% to 2.59%, and GWCC was above 91% for all companies (Table 3). Companies F, A, and E have more nodes with similar connectivity, while D, C, and B have more nodes with dissimilar connectivity. Overall, the network is well-connected, and the GWCC is high for all companies (Table 3.)

**Table 3.** Comparison of six U.S. swine company network parameters. Average and 95% confidence interval of node and edge level network metrics.

| Company | Nodes | Edges | Diameter | Assortativity | Cluster<br>coefficient<br>(10<sup>3</sup>) | GSCC (%) | GWCC (%) |
| --- | --- | --- | --- | --- | --- | --- | --- |
| A | 1,586.82<br>(1,583.44 :<br>1,590.19) | 3,729.07<br>(3,717.67 : 3,740.46) | 11.84<br>(11.61 : 12.07) | 0.35<br>(0.34 : 0.37) | 1.69<br>(1.61 : 1.78) | 0.2<br>(0.19 : 0.2) | 99.2<br>(99.06 : 99.34) |
| B | 279.48<br>(275.42 : 283.54) | 735.84<br>(692.75 : 778.93) | 8.64<br>(8.43 : 8.85) | -0.24<br>(-0.28 : -0.2) | 1.15<br>(0.96 : 1.34) | 0.83<br>(0.64 : 1.02) | 98.83<br>(98.53 : 99.14) |
| C | 171.25<br>(167.91 : 174.59) | 451.92<br>(427.42 : 476.41) | 7.92<br>(6.61 : 9.23) | -0.48<br>(-0.52 : -0.45) | 0.43<br>(0.34 : 0.53) | 3.7<br>(2.04 : 5.36) | 99.62<br>(99.26 : 99.99) |
| D | 838<br>(787.39 : 888.61) | 2,364.45<br>(2,140.01 : 2,588.89) | 11<br>(9.52 : 12.48) | -0.52<br>(-0.55 : -0.48) | 2.08<br>(1.6 : 2.55) | 0.27<br>(0.18 : 0.36) | 97.58<br>(95.56 : 99.61) |
| E | 271.93<br>(268.13 : 275.72) | 451.56<br>(448.46 : 454.65) | 7.7<br>(7.35 : 8.05) | 0.03<br>(-0.01 : 0.06) | 5.63<br>(5.39 : 5.86) | 2.59<br>(2.48 : 2.7) | 91.04<br>(88.47 : 93.61) |
| F | 485.85<br>(475.64 : 496.06) | 1,174.72<br>(1,128.83 : 1,220.6) | 10.28<br>(9.58 : 10.98) | 0.56<br>(0.54 : 0.58) | 3.19<br>(2.92 : 3.46) | 2.31<br>(1.99 : 2.64) | 98.81<br>(98.45 : 99.18) |
| Full<br>network | 2,842.42<br>(2,773.38 :<br>2,911.47) | 6,705.43<br>(6,506.94 : 6,903.92) | 13.3<br>(12.85 : 13.75) | -0.09<br>(-0.11 : -0.08) | 2<br>(1.95 : 2.05) | 0.4<br>(0.36 : 0.43) | 61.58<br>(59.97 : 63.18) |

In Figure 4, we showed county-level in-degree, out-degree, and betweenness. The median in-degree for the three years of movement data was 8 (IQR: 3-15, maximum: 78); out-degree was 8 (IQR: 2-15, maximum: 55); and betweenness median was 36.51 (IQR: 0.4-309, maximum: 7,161.17.)

**Figure 4.**
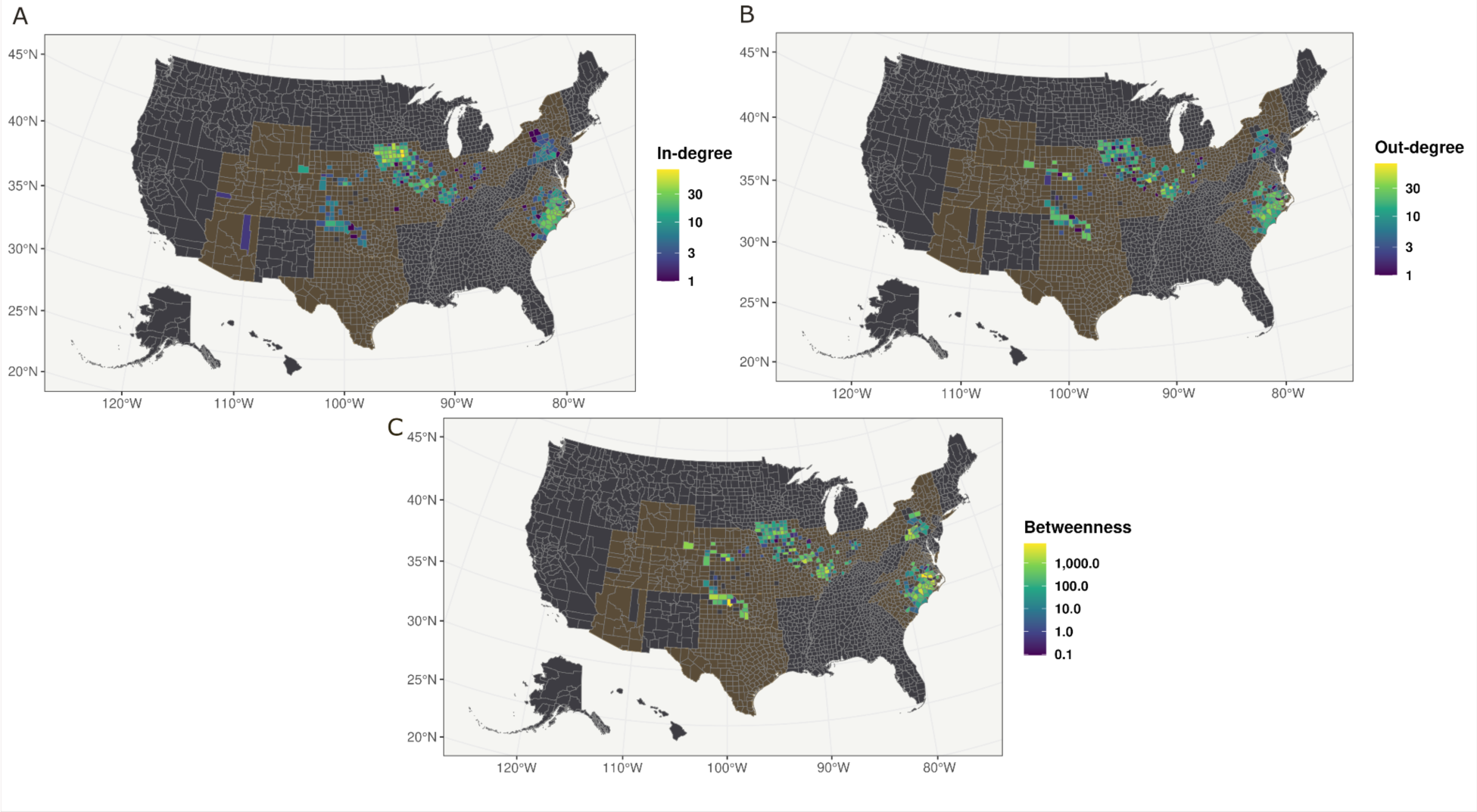
Network measure distribution at the county level. Maps shaded with the static network between county shipments from January 1st, 2020, to January 15th, 2023. States with at least one shipment are highlighted.

### 3.5 Network loyalty analysis

The overall network median in-going loyalty was 0.69 (IQR: 0.60-0.75, maximum: 0.96), and out-going loyalty was 0.52 (IQR 0.36-0.81, maximum: 0.91). Sow and nursery out-going loyalty was the highest, in comparison with wean-to-finisher and finisher loyalty (Figure 5.) Regarding in-going loyalty, finisher, sow, and nursery had similar distributions, with a median of 0.71 (IQR 0.63-0.77, maximum: 0.94), 0.68 (IQR 0.61-0.73, maximum: 0.88), and 0.77 (IQR 0.68-0.82, maximum: 0.96), respectively, and wean-to-finishers and finishers exhibit greater loyalty towards their incoming movements than outgoing movements (Figure 5.) Supplementary Material Figure S13 illustrates loyalty distribution by production type per company, and a descriptive analysis of in-loyalty and out-loyalty values by production type is also shown in Supplementary Material Table S5.

**Figure 5.**
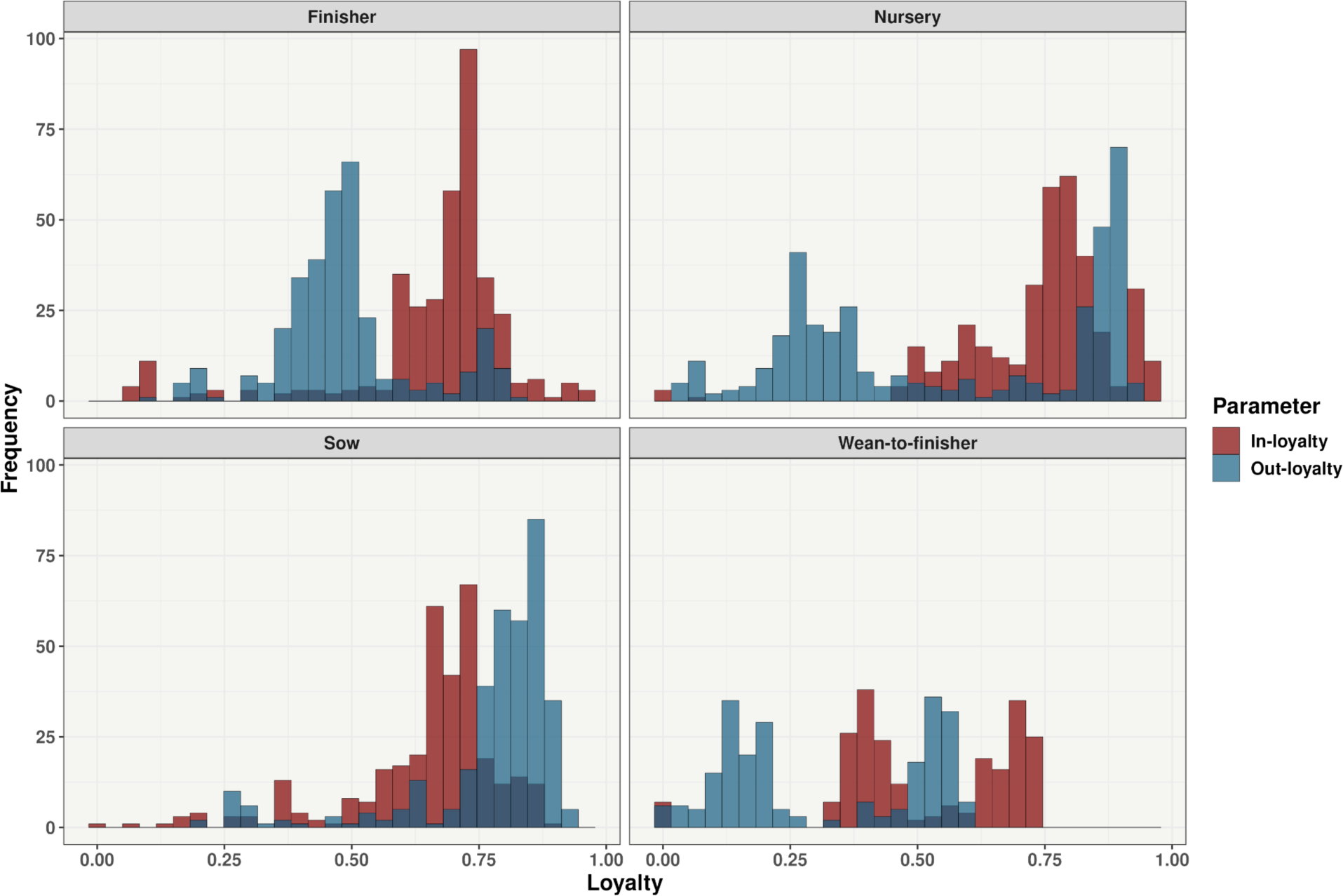
Farm-type loyalty distributions. Histograms are 180 days with weekly time steps by production types. Out-loyalty–whether the production type preserves the same destination premises–is in blue, and in-loyalty–whether the production type maintains the same source premises–is in red.

### 3.6 Dynamic premises-level network degree distribution

Figure 6 shows the in-degree and out-degree distributions, with the out-degree distribution being more skewed than the in-degree. The in-degree distribution peaked at one, with a long tail (Figure 6). The out-degree distribution also peaked at one, with a long tail extending to 30. We also demonstrated that the out-degree distribution followed a power-law distribution with an exponent of α = 3.34 (p-value = 0.75). However, the in-degree distribution did not follow a power-law distribution and only fitted to an α = 4.1 (p-value = 0.083). The Supplementary Material Figure S14 shows the degree distributions for each production type; briefly, out-degree distributions for all production types follow a power-law distribution, while in-degree distributions of finisher and boar stud production types were the only ones that did not fit a power-law distribution.

**Figure 6.**
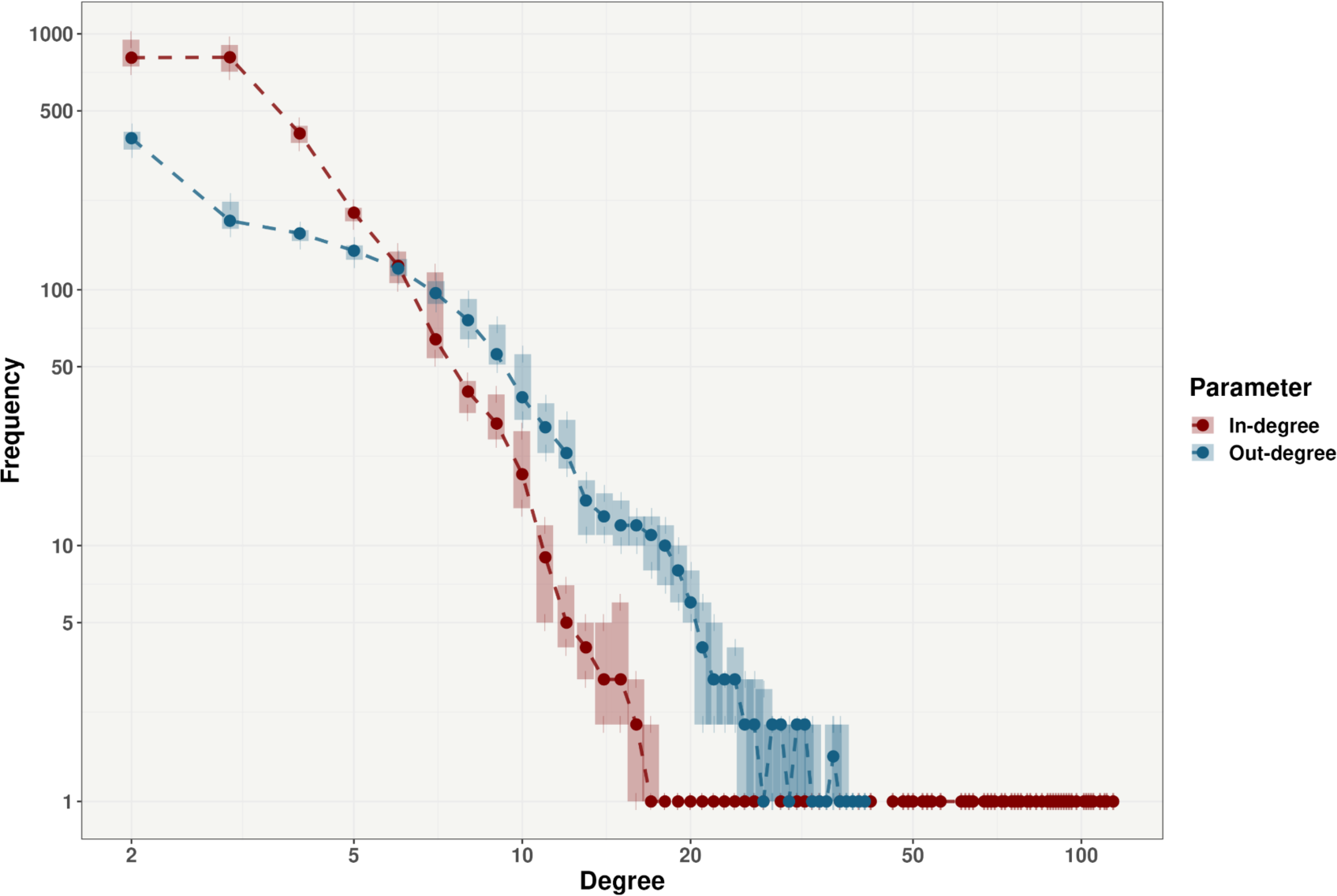
Node degree distribution. In blue, 180-day out-degree intervals, with the median as dots and interquartile ranges as bars. In red are 180-day in-degree intervals, with the median as dots and interquartile ranges as bars. The y-axis and x-axis are on a log10 scale.

### 3.7 Between-premises cumulative contacts as a proxy of outbreak size and effectiveness of network-based targeted percolation

The best network parameters to reduce cascade spread in descending order were degree, betweennesses, cluster coefficient, and PageRank. Cascade reductions were most effective when 25% of nodes ranked by degree were removed, limiting possible spread to 1.23% of premises (IQR: 0.66% -2.38%, maximum: 7.10%). Node removal based on betweennesses was the next most effective, with a median reduction of 1.70% (IQR: 0.87%-3.21%, maximum: 20.42%). Cluster coefficient and PageRank reduced cascade to a median of 4.87% (IQR:2.15%-4.96%, maximum: 53.63%) and 8.09% (IQR: 2.57%-16.64%, maximum: 56.40%), respectively. As expected, random node removal had the most negligible effect, reducing cascades to a median of 12.01% (IQR: 3.42%-26.34%, maximum: 65.68%). We noted similar overall patterns when percolation was applied to each company’s network (Supplementary Material Figure S15.)

**Figure 7.**
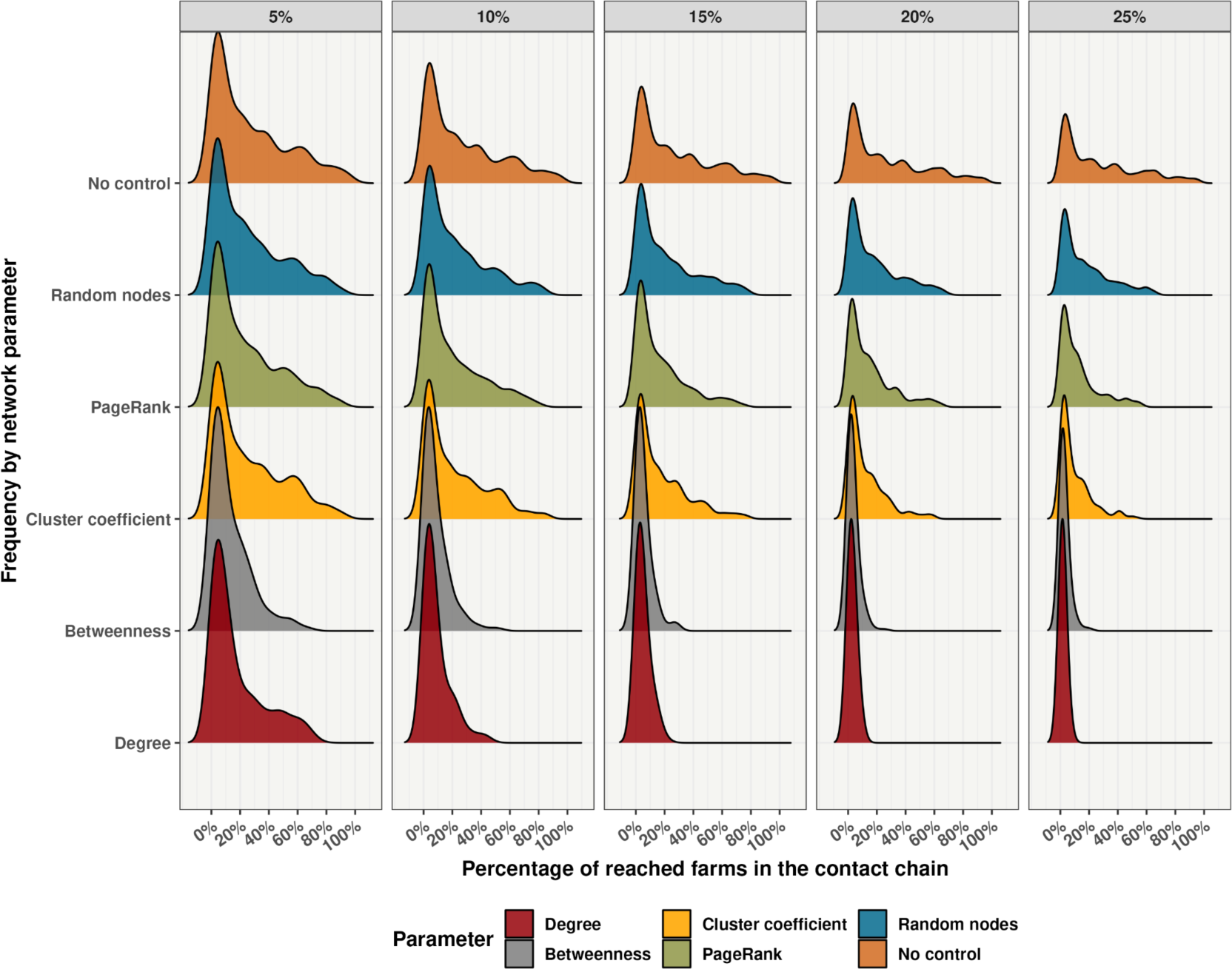
Density distribution of network-based node percolation removal. The ridgeline plot in the top panel shows the percentage of removed premises (5%, 10%, 15%, 20%, and 25%) for network parameters (y-axis). The x-axis shows the percentage of premises within cascades after node removal.

## 4. Discussion

In this work, we examined the patterns of between-premise pig movements of six U.S. commercial swine production companies with premises within 20 U.S. states. While most pig shipments were intrastate, with a median of 52 km between the source and destination premises, we demonstrated the presence of long-distance, interstate shipments occurring over distances greater than 1,000 km. We showed that the highest number of intrastate pig shipments among our dataset were in North Carolina, Iowa, and Oklahoma. Shipment patterns significantly differed across companies and production types, with the most significant differences observed in commingling and between premises movement loyalty. The median number of pig shipments of different sources mixing at destination premises varied between two and four. In-going loyalty was at 69%; thus, within 180 days, 69% of movements came from the same premises, and out-going was 52%, indicating lower loyalty for outgoing movements. Overall, sow premises had higher out-going loyalty, implying that piglets often are weaned into the same downstream farms. In contrast, wean-to-finisher, had the lowest in-going loyalty, thus receiving pigs from a wide range of farms. The out-degree distribution fitted a power law distribution, exhibiting scale-free topology, which suggests the presence of highly connected premises (hubs) with the capacity to drive super-spread disease events. On the other hand, such scale-free topology is ideal for network-based target control tactics, as shown by the network-based targeted percolation approach, in which targeting farms by degree was highly effective in reducing potential epidemics.

Long-distance disease spread events have been directly associated with the shipment of pigs (Buhnerkempe et al., 2014; Fèvre et al., 2006; Galvis et al., 2022b; Relun et al., 2016; Sykes et al., 2023). An in-depth understanding of national movement patterns is a key element for epidemic control in highly integrated swine production systems. In the same vein, examining interstate and intrastate pig shipment data is crucial to informing national disease response plans, and developing novel tactics against endemic and emerging diseases (Cardenas et al., 2022; Hammami et al., 2022b; Puspitarani et al., 2023; Sykes et al., 2023). The median distance of pig shipment was 74 km, similar to what has been shown in other regional studies with shipment distances between 10 and 50 km (Kinsley et al., 2019). Moon et al., 2019 used Iowa State census data and estimated that most shipments were more likely to occur at 20 km. While we demonstrated that most movements occurred over short distances and were intrastate movements, we noted significant differences in shipment distances among companies (Supplementary Material Figures S9 and S10). Most long-distance shipments were related to boar and replacement gilt shipments (Figure 3), similar to what has been described elsewhere (Blair and Lowe, 2019; Kinsley et al., 2019; Subharat et al., 2022). In addition, considering the shortest roadways, most movements were within North Carolina, 3,446 (25%) roads. We also showed a limited set of roadways, 382 (1.49%), connecting North Carolina to Pennsylvania, Ohio, Iowa, and Oklahoma. Ultimately, mapping road pathways may be used to develop disease control tactics and aid in logistical planning by identifying less congested routes, which can be important for transporting highly valuable animals, such as breeding pigs and boars.

The dynamic network analysis revealed that the GSCC ranged from 0.2% to 2.59%, while the GWCC varied from 91.04% to 99.62%. These results highlight the extreme levels of verticalization of North American commercial swine production, as sows typically send weaned pigs to the same nurseries or wean-to-finisher premises and then to a wider range of finishers (Kinsley et al., 2019; Machado et al., 2019; Passafaro et al., 2020), similar patterns have also been described in Austria (Puspitarani et al., 2023) and Brazil (Cardenas et al., 2022, 2021). Time-independent, connected component sizes provide an overall estimation of the lower and upper bounds of epidemic sizes (Kao et al., 2006; Omondi et al., 2021); for the U.S., the upper bound estimates above 90% of premises are at risk of infection.

Low out-going loyalty was significant in finisher, nursery, and wean-to-finisher across all companies (Table 2 and Supplementary Material Figure S13). Such low loyalty, when associated with large out-going contact chains, may potentially increase their risk of becoming super-spreaders. Similarly, premises with low in-going loyalty and high commingling are expected to act as super-receivers with a higher risk of infection (Acosta et al., 2023). In the present study, we identified extreme examples of finishers and nurseries receiving or sending animals to up to 144 premises. At the same time, such premises were less than ten distinct sites; these farms can be classified as super receivers and spreaders (Table 2 and Supplementary Material Figure S13) for which the implementation of movement restrictions would result in the immediate stop of the spread of the disease. On the other hand, premises with higher loyalty values and lower commingling are expected to remain uninfected for longer periods (Makau et al., 2021; Schulz et al., 2017). Our study found similar loyalty distributions as Makau et al., 2021, which reported a loyalty of 0.63. However, when we compared our results to the German swine network, we found that breeding and multiplier premises had a higher out-loyalty, ranging from 0.49 to 0.67 (Schulz et al., 2017). This suggests that there is significant variation in the topology of the swine production network across countries, which is likely to have a direct effect on the effectiveness of movement restriction strategies to control the spread of infectious diseases.

Out-degree distribution followed a scale-free distribution; thus, most premises are poorly connected, and a few have a very high number of connections. Such network property is rare among natural systems (Broido and Clauset, 2019) and has been noted by other swine network studies (Passafaro et al., 2020; Relun et al., 2016). We demonstrated that in-degree exponents were higher than out-degree exponents, suggesting greater heterogeneity for recipient premises than for source premises. We have also shown that degree distribution was similar among production types. Still, for wean-to-finisher premises, the in-degree distribution density between five and ten degrees was more frequent when compared with the out-degree distribution (Supplementary Material Figure S14); therefore, wean-to-finisher premises are more likely to be “super-receivers” and nurseries and sow farms “super-spreaders.” Given the degree distributions observed in this study, network-based control and prevention strategies will significantly impact containing movement-related dissemination events. Indeed, we demonstrated that a median of 19.1% of premises would likely be infected if no movement restriction control actions were implemented; however, when targeting 25% of the premises based on degree and betweenness, we were able to reduce the spread to 1.23%, and 1.7% of infected premises, respectively. This result aligns with other movement restriction methods in which degree and betweenness efficiently decrease the transmission chain in a temporal network or fragment the large connected components in the network (Cardenas et al., 2022; Kinsley et al., 2019; Lentz et al., 2016; Passafaro et al., 2020, 2020). The spread cascade model proposed in this study could, therefore, be used as an additional easy-to-calculate parameter when planning surveillance activities and developing disease contingency programs.

In the U.S., between-state livestock shipment data is generally collected via certificates of veterinary inspection (CVIs) (Gorsich et al., 2019), while within-state shipment data are collected routinely by industry operators but may not be required to be shared with animal health officials (Cabezas et al., 2021). Previous studies describing U.S. swine contact networks have used interstate certificates permitting the movement of animals (Cabezas et al., 2021; Gorsich et al., 2019; Passafaro et al., 2020). However, due to the lack of large-scale, real, between-farm swine movement data, Passafaro et al., 2020, adapted a Bayesian Markov chain Monte Carlo model used to simulate cattle shipment networks (Lindström et al., 2013) to simulate swine shipment networks. These simulated networks included predictions of shipment sizes, number of nodes, and edges and were used to generate maps of movement distributions at the county level. Here, we use our between-premises movement data to provide a side-by-side comparison between real data and simulated results (Passafaro et al., 2020). Our comparison indicated significant differences in the in-degree, out-degree, betweenness, and outgoing number of pigs values between real and simulated data (Kruskal-Wallis Rank Sum Test, p-value <0.05). The simulated data overestimated the in-degree and out-degree by 20 to 22 degrees, respectively, and underestimated the total degree of 43 and 44 counties of 100 counties (Supplementary Material Figure S16-S18). Ultimately, we argue that while simulated networks may be helpful in the absence of true movement data, the results fail to represent the complexity of real-world commercial swine network topology. Therefore, we highlight the relevance and need for real movement data for better disease control preparedness and suggest caution in using simulated datasets.

## 5. Limitations and further remarks

While we described the pig movements of commercial swine-producing companies, our results need to be interpreted cautiously as it does not include all commercial swine-producing companies within the 20 U.S. states we presented. We used the premises’ unique identification to reconstruct the contact network; we acknowledge that 10.16% of premises had more than one production phase in the same premises (e.g., farrow to the finisher). In those cases, we could not identify the production phase receiving the movements; thus, commingling results could have been impacted. Another limitation relates to the node removal results, which are a pure network-based approach used to identify target farms based solely on their pig movement network patterns; this approach does not consider other indirect contacts, such as the movement of vehicles and/or trailers (Sykes et al., 2023; Galvis et al., 2022a, 2021); thus, results should be interpreted with caution. Additionally, the results of node removal do not consider any disease detection rate variations, thus, we remark that such strategy effectiveness is impacted by several factors, for instance if infected farms are not detected immediately node-level control actions is severely reduced(Tago et al., 2016).

Overall, this study is the first to use large real pig movement data from several commercially integrated companies. This offers an opportunity to explore intrastate pig movement patterns, which remain mostly disaggregated in the U.S., while also revealing the volume of interstate movements associated with these swine companies. Our results could help policymakers target national disease control and surveillance plans while providing the swine industry with a comprehensive view of premises-to-premises movement patterns.

## 6. Conclusion

The results of this study provide pivotal information to understand the topology of the network structure of interstate and intrastate pig of six U.S. commercial swine production companies with premises within 20 U.S. states. We showed that most movements were within 74 km of the origin premises, which is relevant information for disease control and prevention. We quantified the mixing of different animal sources (commingling) and loyalty of premises to provide practical information to inform risk-based health interventions; premises with low loyalty and high commingling values, i.e., finisher farms, are at greater risk of becoming infected via the movement of pigs. Our spread cascade model, based on out-going contact chains, was a useful approach to prove that targeting farms by degree and betweenness when implementing control actions (i.e., vaccination, increased quarantine, and improved hygiene protocol) would be effective in restricting disease spread, which helps to improve the efficiency and efficacy of disease surveillance and management of an infectious disease.

## Acknowledgments

The authors would like to thank the participating companies and veterinarians for their insightful comments and engaging discussions.

## Authors’ contributions

NCC and GM conceived and designed the study. NCC, AV, and FS conducted data processing, cleaning, and analysis. All authors wrote and edited the manuscript. All authors discussed the results and critically reviewed the manuscript. GM secured the funding.

## Ethical statement

The authors confirm the ethical policies of the journal, as noted on the journal’s author guidelines page. Since this work did not involve animal sampling or questionnaire data collection by the researchers, there was no need for ethics permits.

## Data Availability Statement

The data that support the findings of this study are not publicly available and are protected by confidential agreements; therefore, they are not available.

