## supplementary material for "Analyzing the intrastate and interstate swine movement network in the United States"

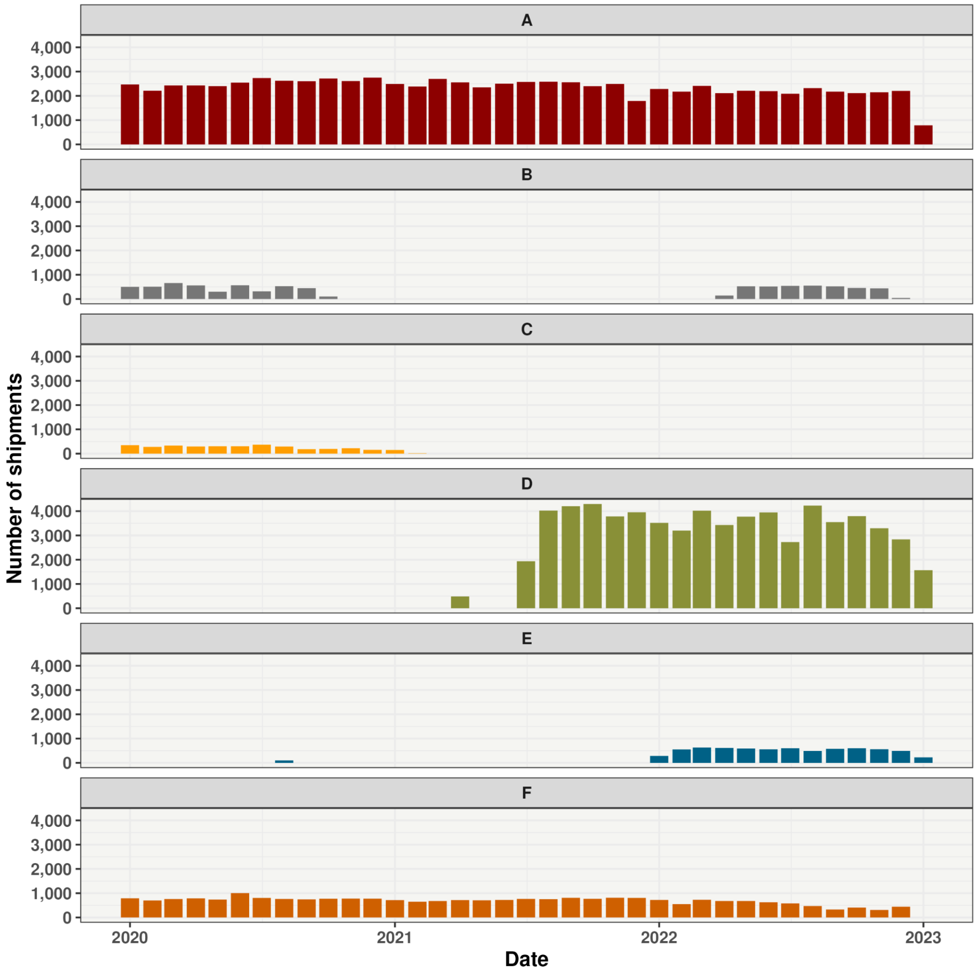


**Supplementary Figure S1.** The monthly number of between-farm shipments by swine production companies.

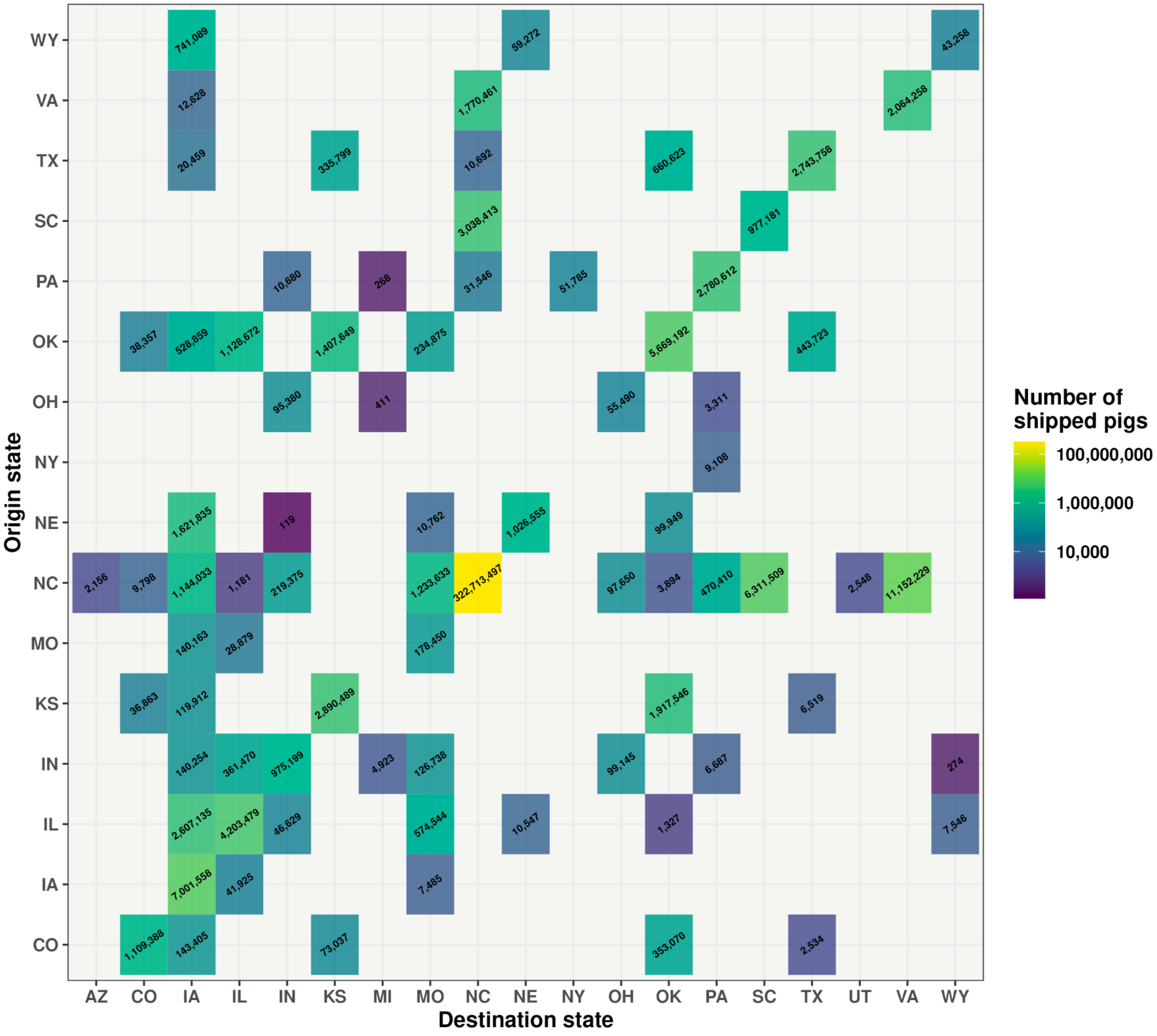


**Supplementary Figure S2.** The number of interstate and intrastate pigs shipped among U.S. states.

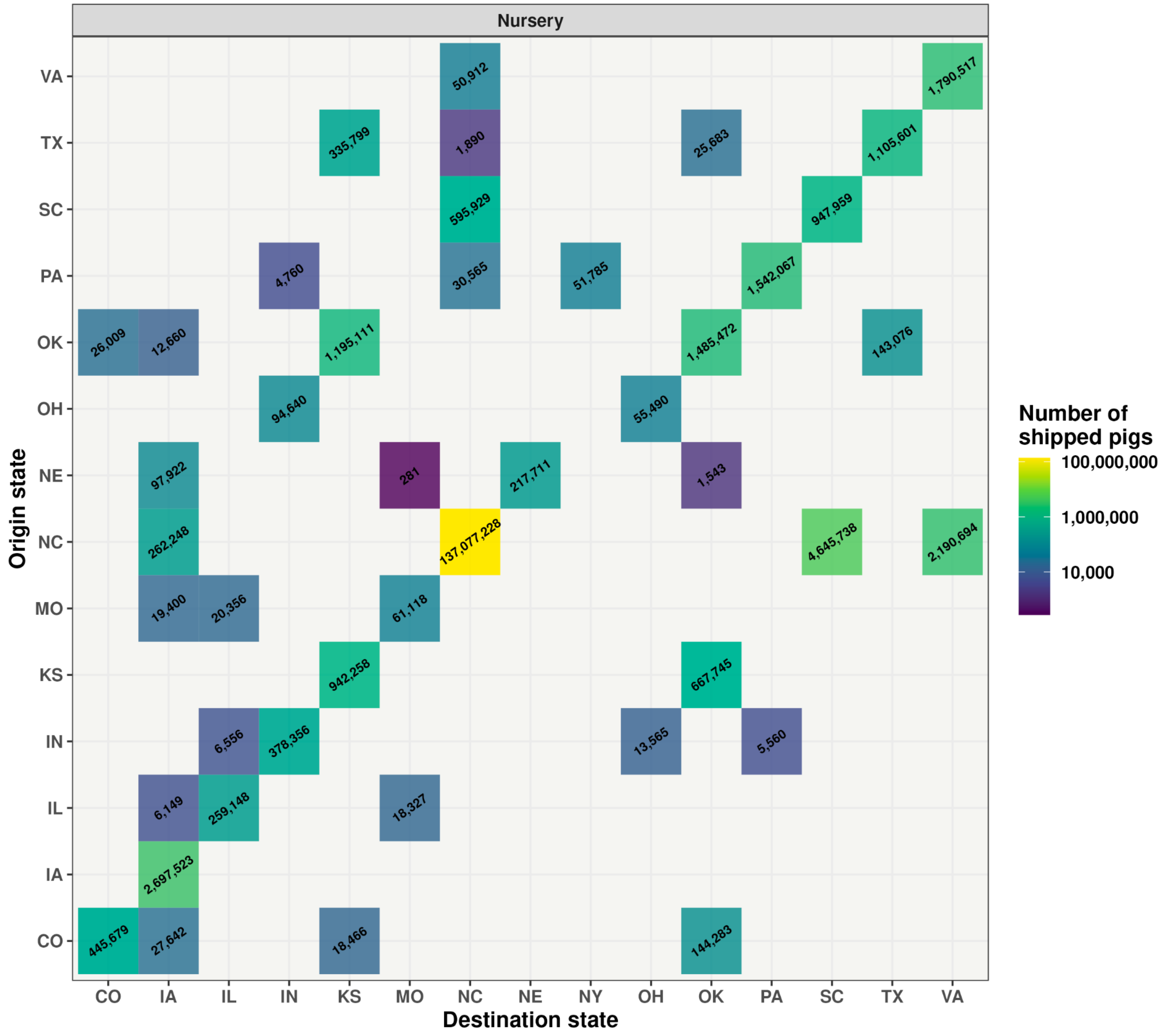


**Supplementary Figure S3.** The number of interstate and intrastate nursery pigs shipped among U.S. states.

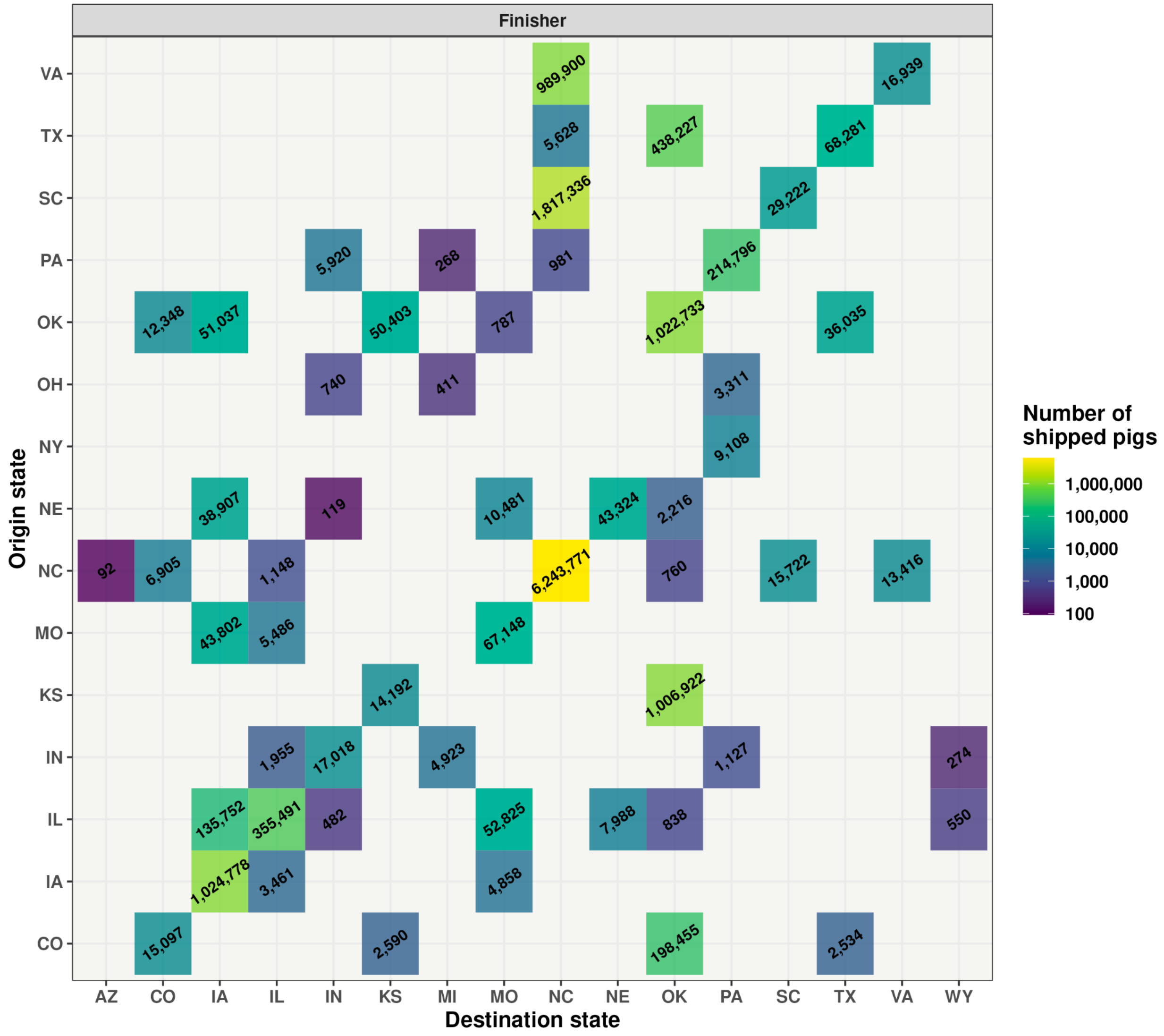


**Supplementary Figure S4.** The number of interstate and intrastate finisher pigs shipped among U.S. states.

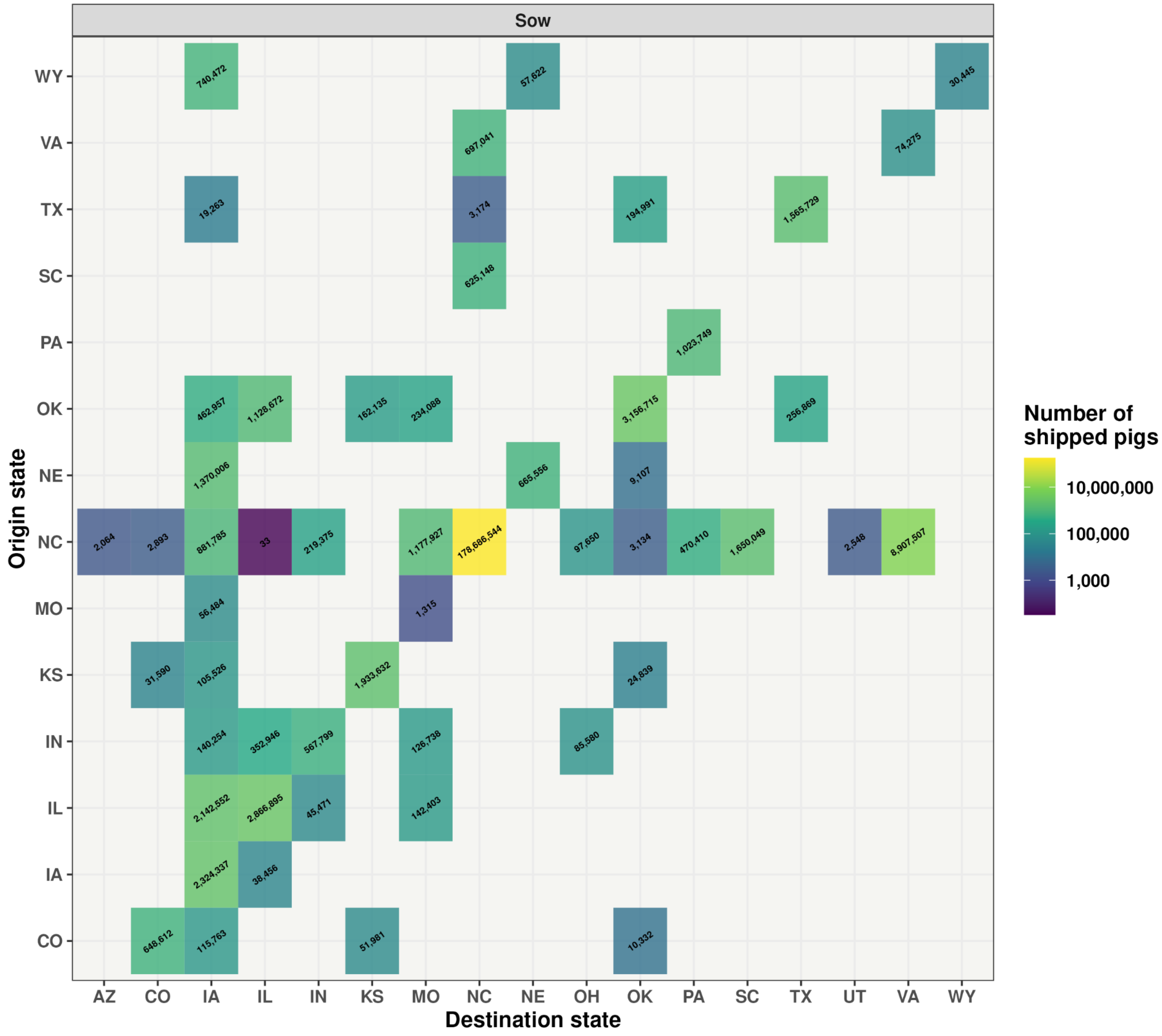


**Supplementary Figure S5.** The number of interstate and intrastate sow pigs shipped among U.S. states.

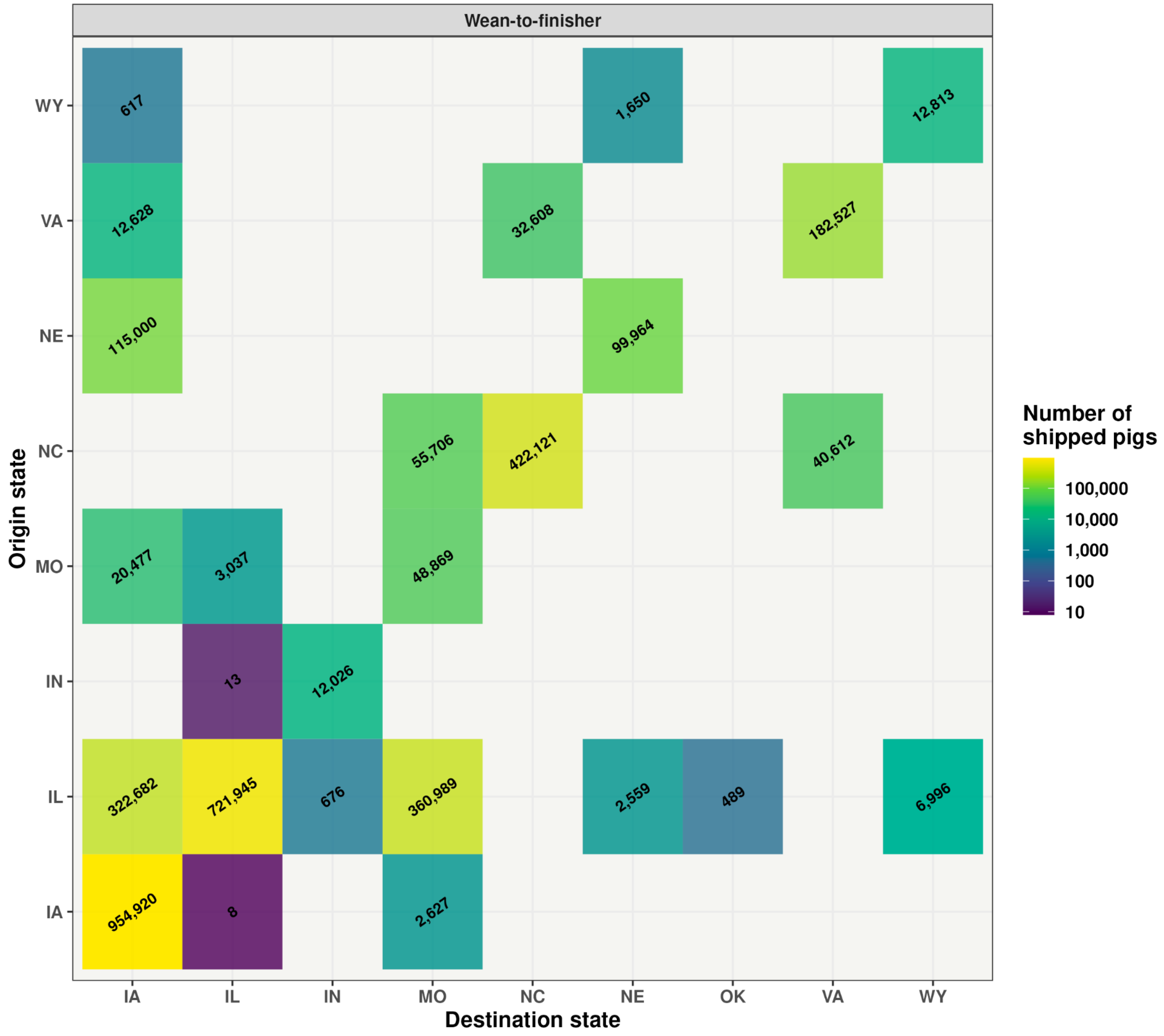


**Supplementary Figure S6.** The number of wean-to-finisher pigs shipped among U.S. states.

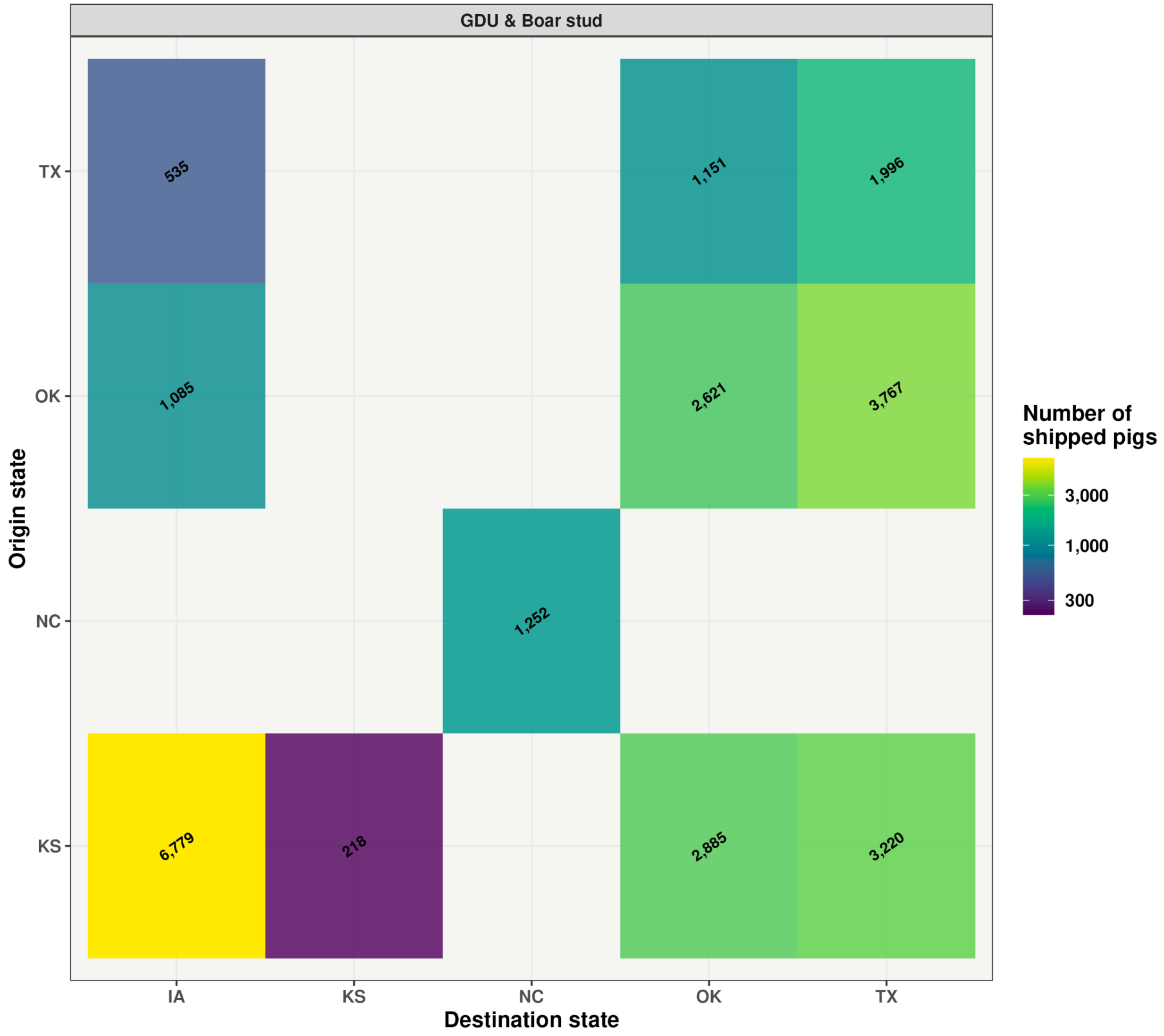


**Supplementary Figure S7.** The number of gilt development unit (GDU) and board stud pigs shipped among U.S. states.

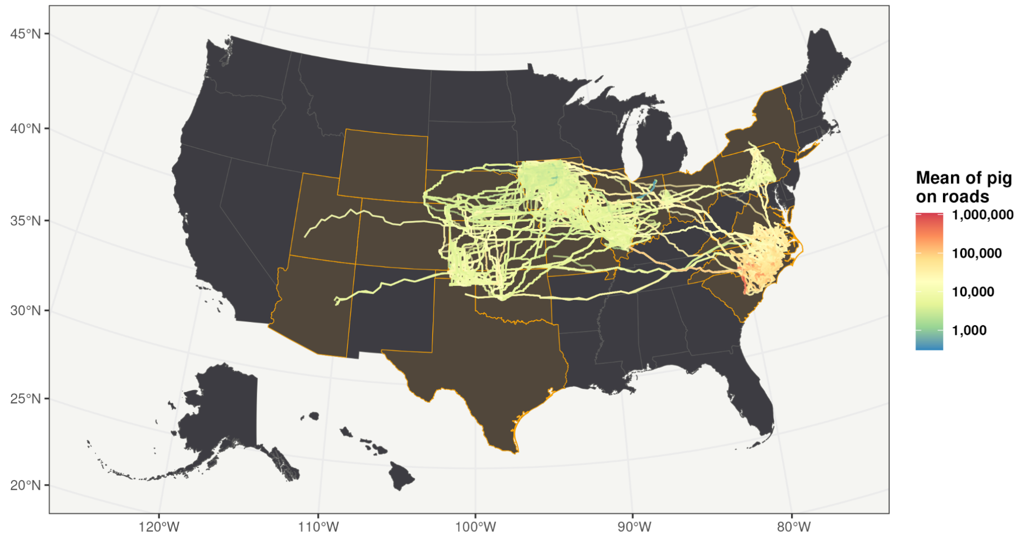


**Supplementary Figure S8.** Map of the average number of pigs shipped by road. The lines represent the routes from the origin and destination premises, and the color represents the mean number of shipped pigs.

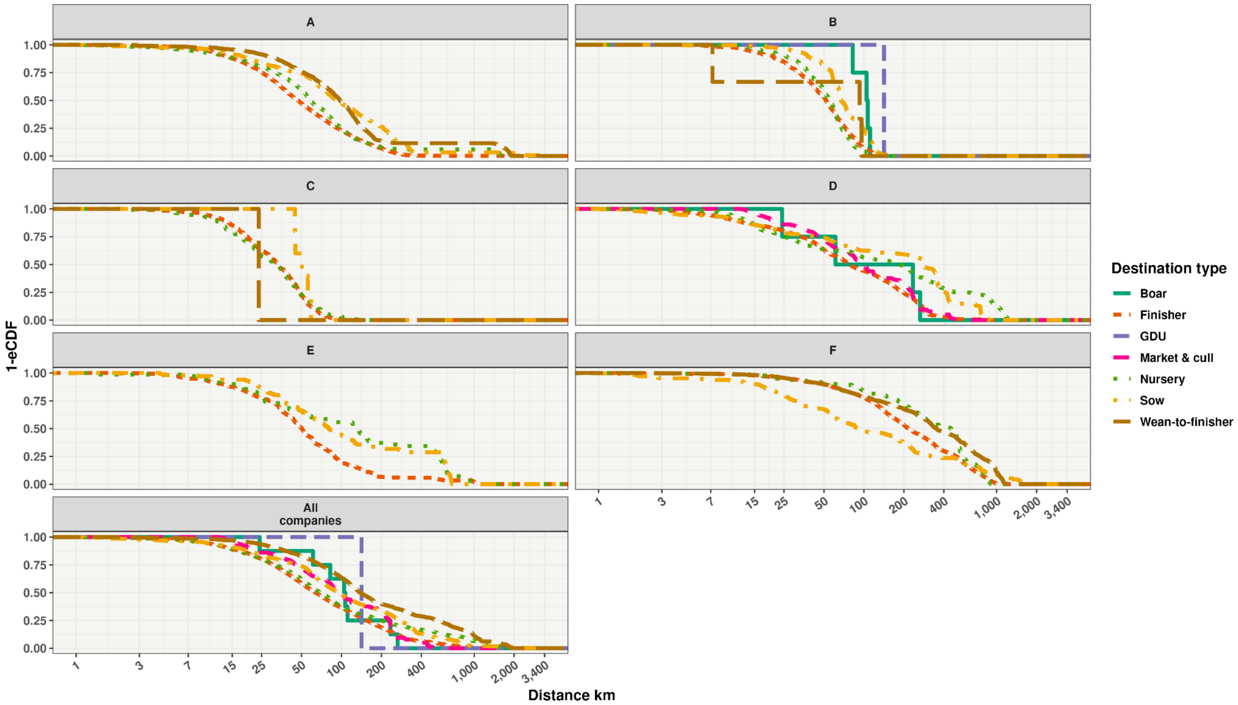


**Supplementary Figure S9.** Distribution of road distances when considering the origin type.

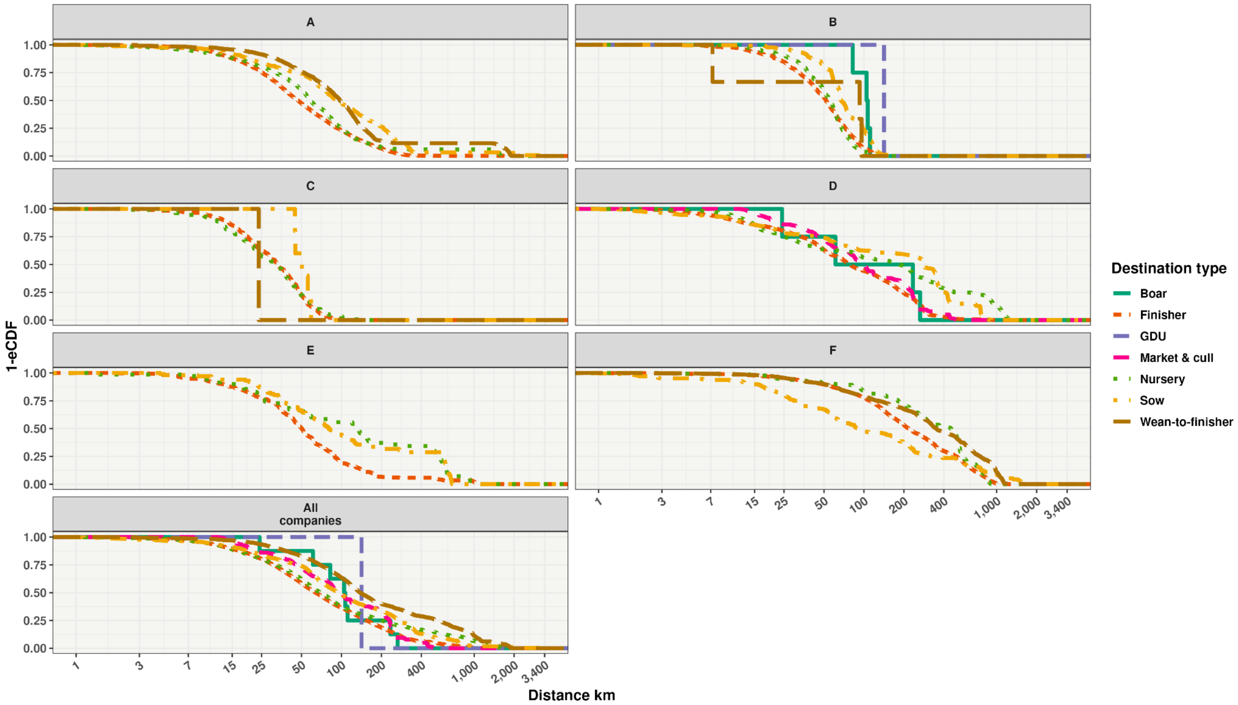


**Supplementary Figure S10.** Distribution of road distances when considering the destination type.

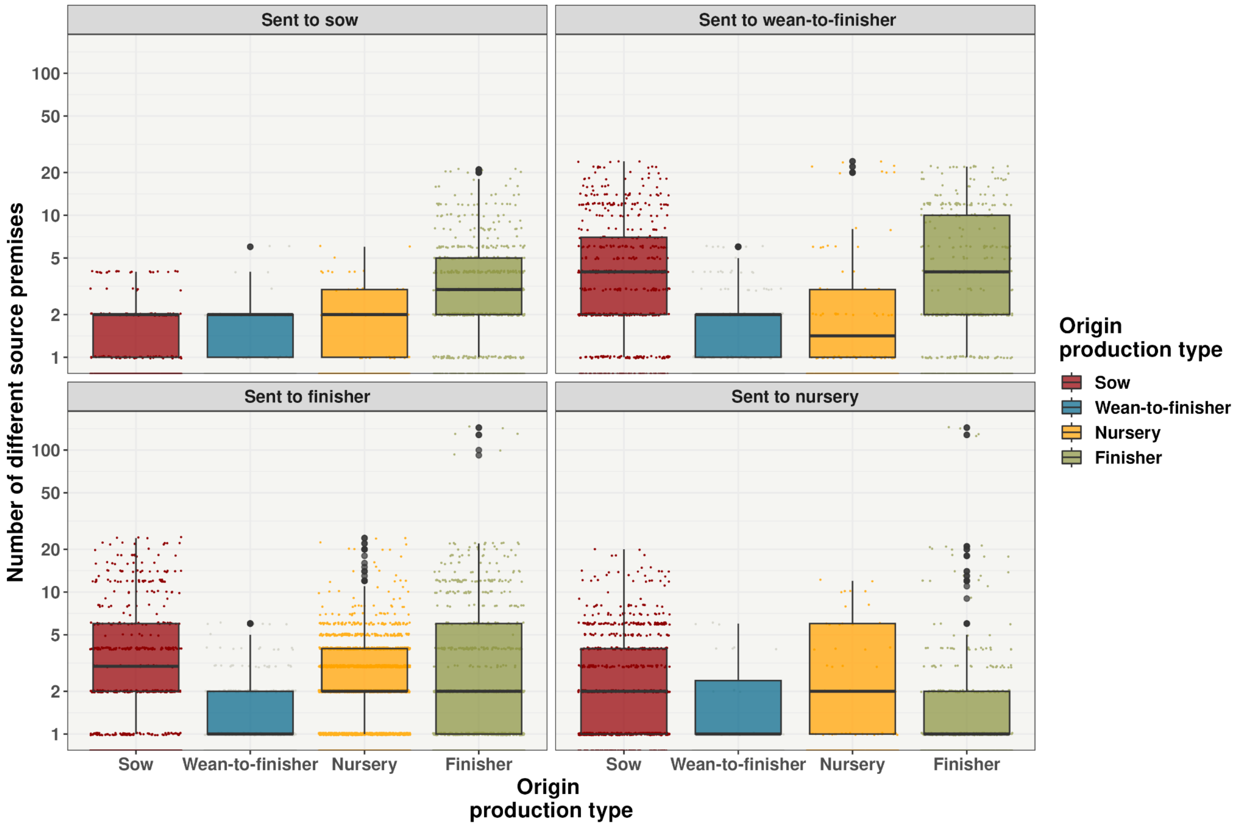


**Supplementary Figure S11.** Commingling at premises level. The x-axis displays the origin production type while each panel indicates the type of destination premises, and the y-axis shows the distribution of the number of in-going commercial partners.

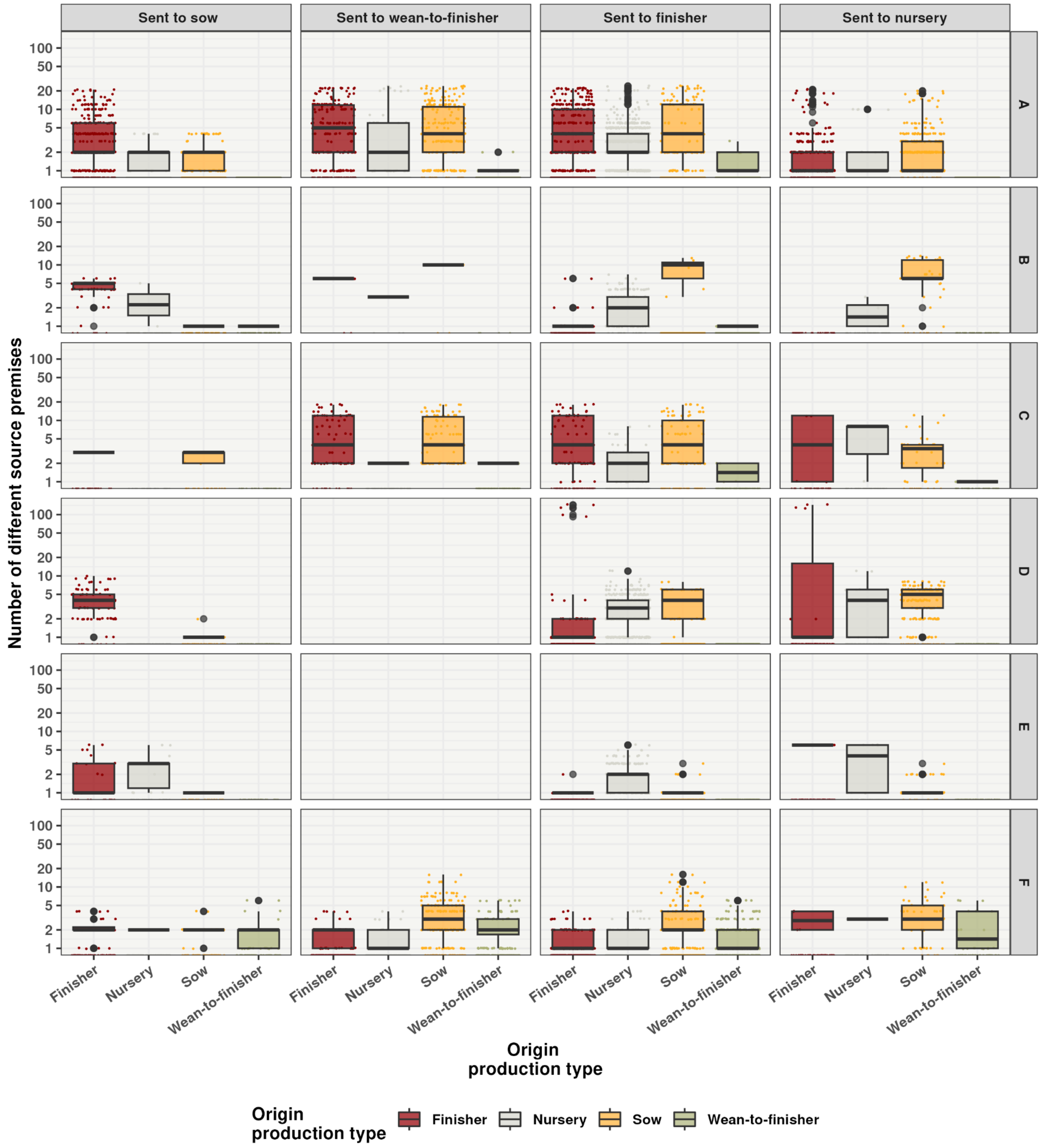


**Supplementary Figure S12.** Premises source mixing (commingling) by company.

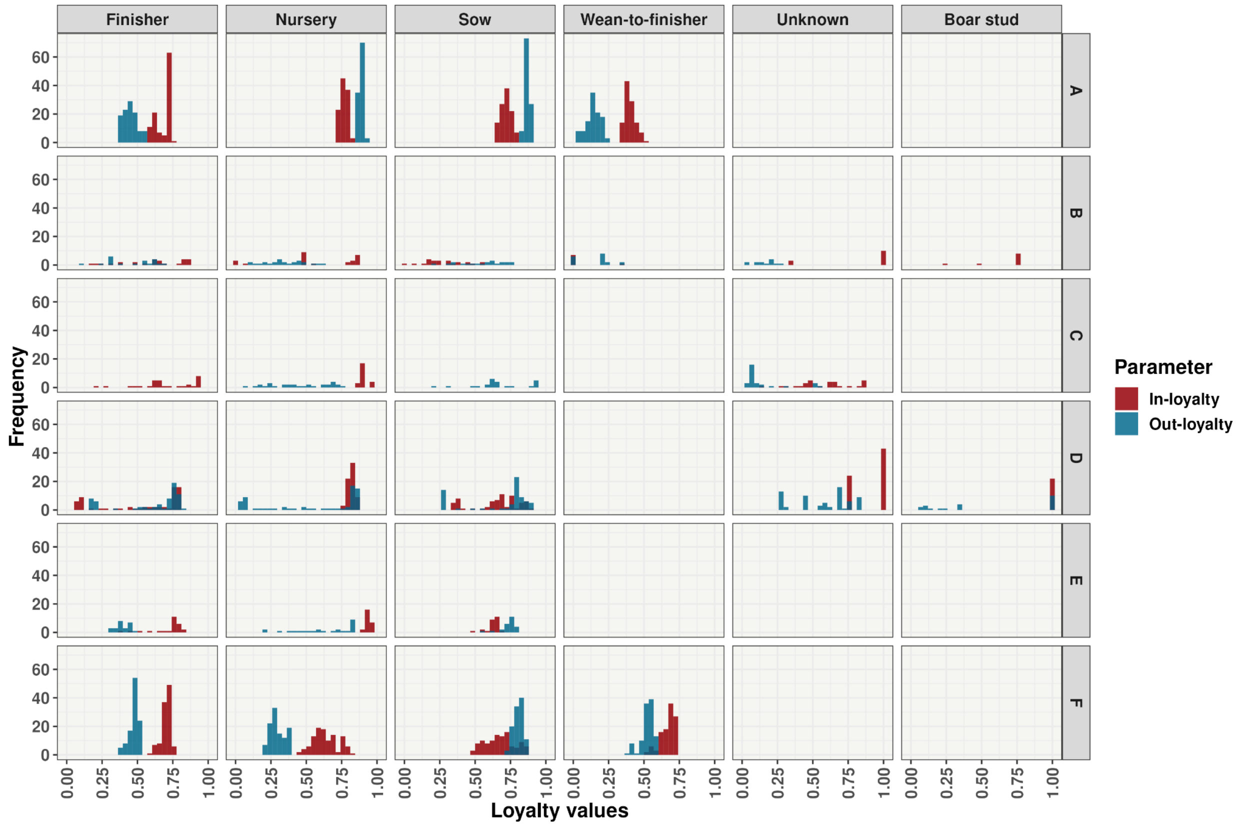


**Supplementary Figure S13.** A 180-day histogram with weekly time steps is presented for the production types of all production types and by company. Out-loyalty is depicted as red-colored hashes, and in-loyalty is shown as blue-colored dots.

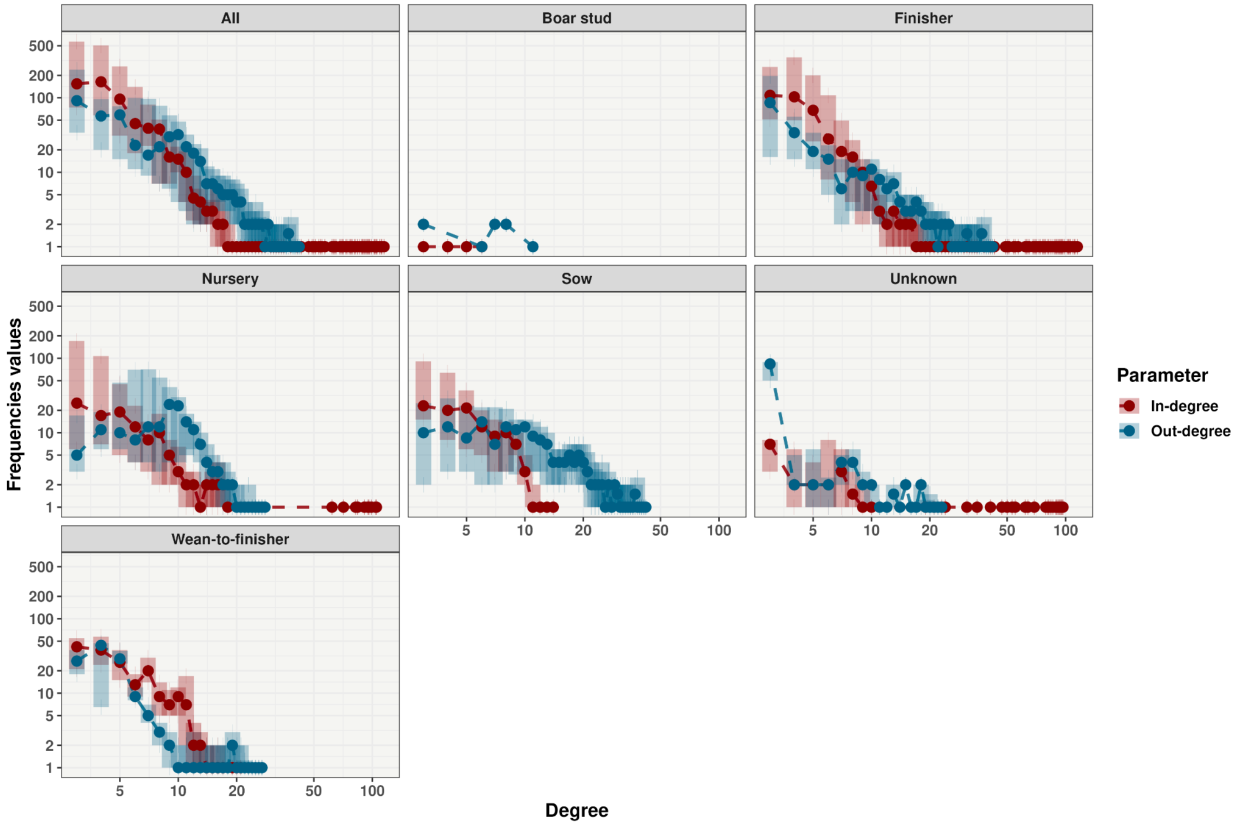


**Supplementary Figure S14.** Degree distribution by production type. In blue, we show 180-day out-degree intervals, with the median as dots and interquartile ranges as bars. In red, 180-day in-degree intervals, with the median shown as dots and interquartile ranges as bars. Both axes are on a base 10 logarithmic scale.

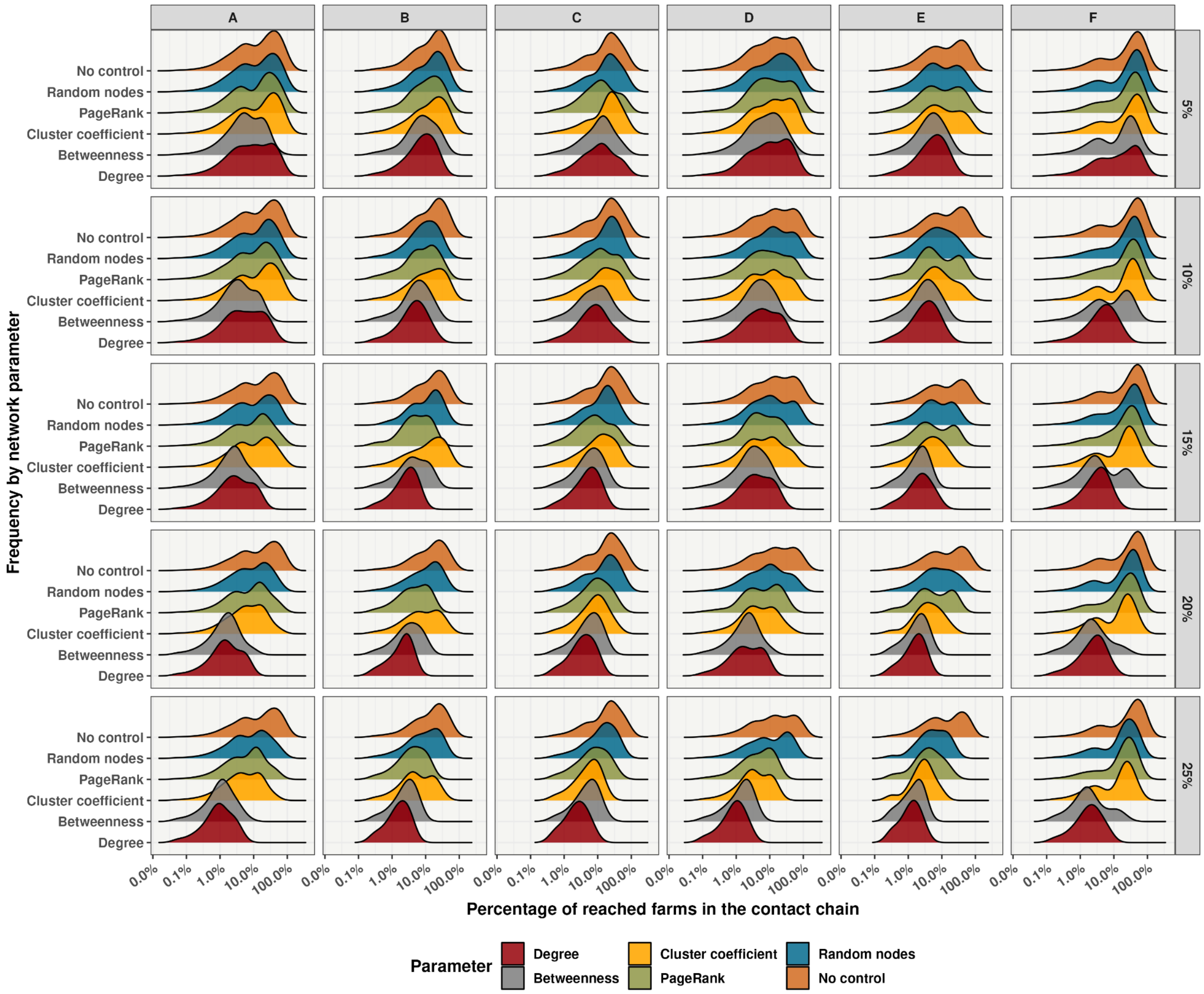


**Supplementary Figure S15.** Spread cascade chain after node removal by company. A Ridgeline plot shows the distribution of a numeric value for network parameters. Distribution was represented using density plots, all aligned to the same horizontal scale and presented with a slight overlap. The y-axis represents the frequency of the out-going contact chain for each premise by the production company (top panel) and by the percentage of nodes removed according to a specific node parameter (right panel).

**Comparison between real between-farm movement data (this work) and simulated movement data (Passafaro et al., 2020)**

We aimed to compare the real data with simulated data (Passafaro et al., 2020) by analyzing in-degree, out-degree, and betweenness and outgoing movements. Briefly, the simulated data includes 3,079,839 records and 250 simulation repeats; for in-depth details about the methods, read (Passafaro et al., 2020). Simulated data utilized the county of origin and destination to reconstruct the contact networks. The comparisons were restricted to results of movements among North Carolina counties due to the completeness of the data available. We constructed a directed network using the counties as nodes and the simulated movements as edges to calculate in-degree, out-degree, betweennesses, and outgoing movements. Since there are 250 different replication results in the (Passafaro et al., 2020) study, we obtained 250 different networks; thus, to obtain a unique result by county, medians of the in-degree, out-degree, betweennesses, and outgoing movements were calculated. This data is hereinafter referred to as "simulated" data. To compare the real data with the simulated results, we reconstructed the animal movement network at the county level by using counties as nodes and the animal movement records as edges; this data is hereinafter referred to as "real data" data. We quantified the difference between the in-degree and out-degree values observed in the simulated and the real data. Results are shown in maps plotted at the county level (Supplementary Figure S16 to S18). Figure S16 shows the over-smoothing of the simulated movement data to counties with no known commercial farm locations and elevated degrees and betweenness in counties to highlight pig populations. In addition, in Figure S17, we provide the difference between simulated and real data. Positive values show where simulated data (Passafaro et al., 2020) overestimated in-degree and or out-degree, while negative values showed counties in which the simulated approach (Passafaro et al.,

2020) underestimated movement. In Figure S18, we overlap the movement frequencies of real and simulated data. We show that simulated data can match low values, but does not perform well for countries with large degree and betweenness values. Simulated data does better matching the number of outgoing than ingoing movements

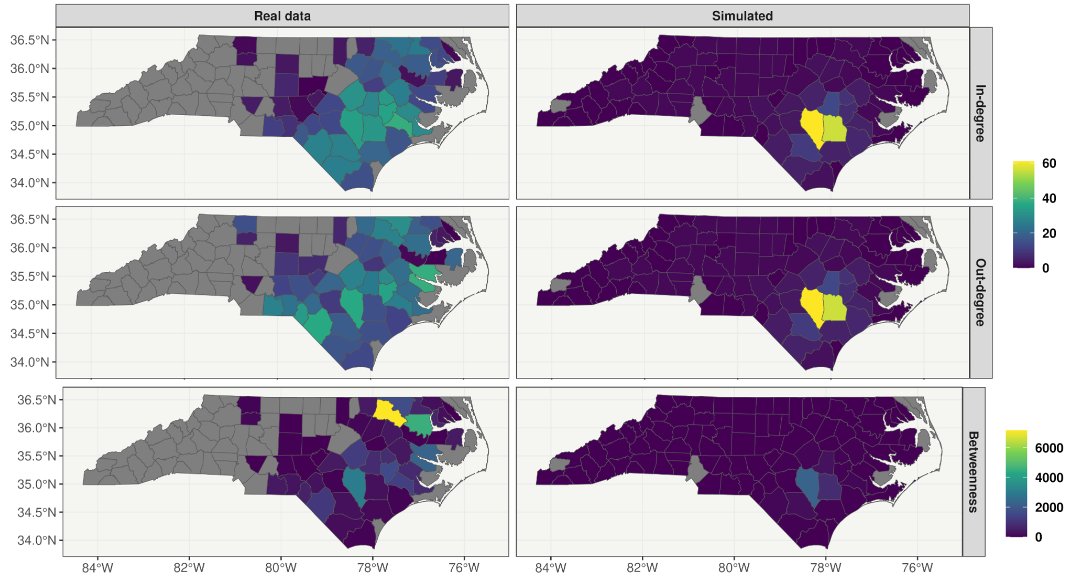


**Supplementary Figure S16.** Maps of North Carolina represent the values of in-degree, out-degree, and betweennesses, simulated data (Passafaro et al., 2020), and real data.

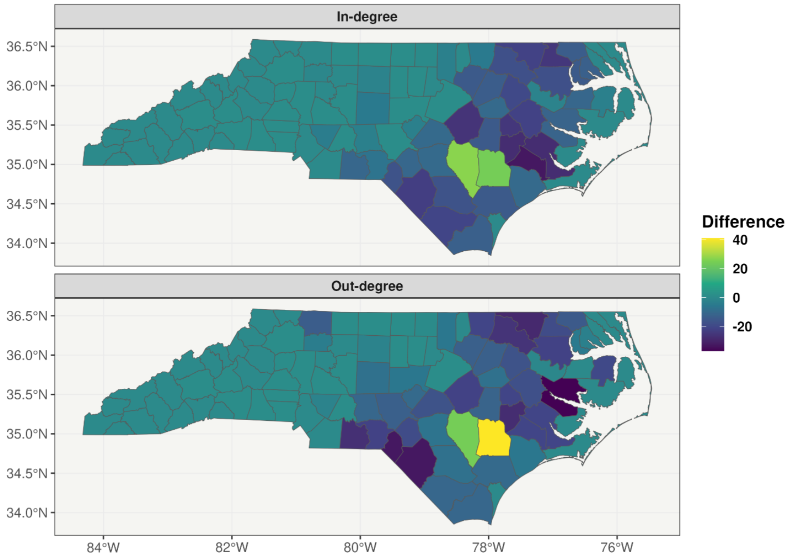

**Supplementary Figure S17.** Difference between the observed in-degree and out-degree between real data and simulated data (Passafaro et al., 2020). Positive values are counties in which the simulated data overestimated the degree compared to the real data. Similarly, negative values represent underestimation.

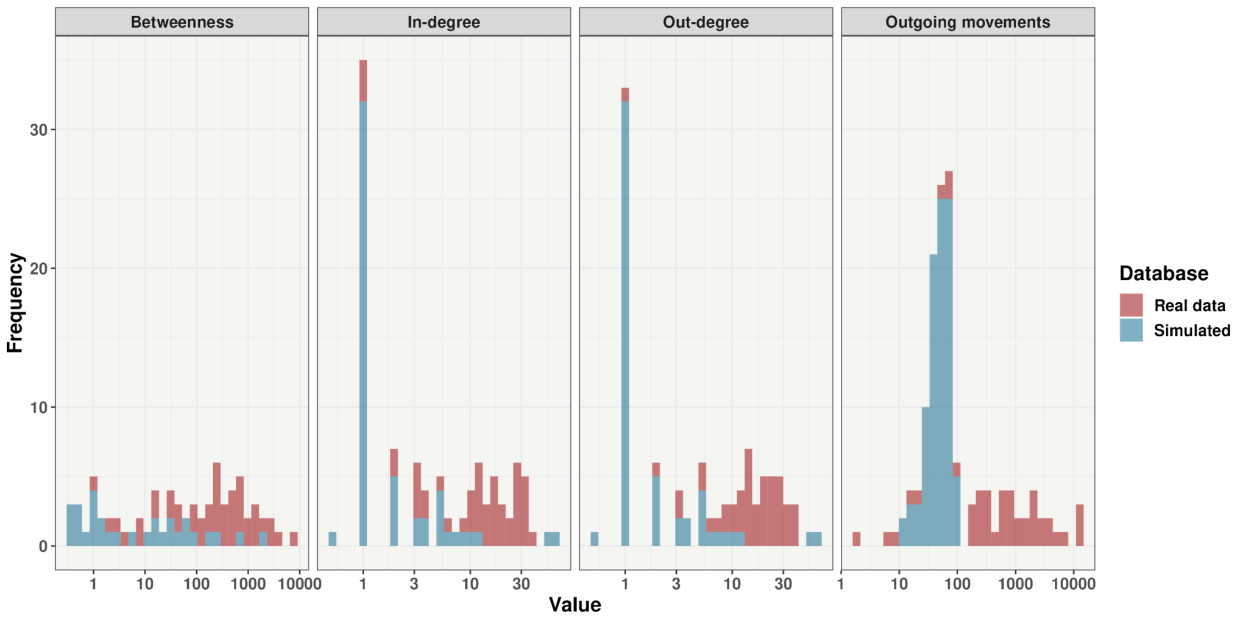


**Supplementary Figure S18.** Histogram of the frequency values of betweenness, in-degree, out-degree, and outgoing number of movements observed in the simulated data (Passafaro et al., 2020) and real data.

**Supplementary Table S1.** Description of network analysis terminology and parameters

| Parameter | Definition | Reference |
|---|---|---|
| Nodes | The unit of interest in network analysis. In the case of this work being the individual premises. | (Wasserman and Faust, 1994) |
| Edge | Link between two nodes in the network. In the case of this paper being the pig movements between two premises. | |
| Degree (k) | Number of unique contacts to and from a specific node (e.g., premises location). When the direction is considered, the in-going and out-going contacts are separated: out-degree is the number of contacts originating from a specific node, and in-degree is the number of contacts coming into a specific node. | |
| Giant weakly connected component (GWCC) | Proportion of nodes that are connected in the largest component when the directionality of movement is ignored. | (Wasserman and Faust, 1994) |
| Giant strongly connected component (GSCC). | The proportion of the nodes that are connected in the largest component when directionality of movement is considered. The percentage of nodes in GSCC and the number of nodes in GWCC complement each other, as the first informs the risk level of the network structure and the second the maximum number of holdings at risk of direct transmission if a disease is found | (Wasserman and Faust, 1994) |
| Network diameter | The length of the shortest path between the most distanced nodes in the network, given that the shortest path actually exists. | (Wasserman and Faust, 1994) |

| | | |
|---|---|---|
| Assortativity | Measures the tendency of nodes of a given degree to be linked to nodes with similar (assortative mixing) or different degrees (disassortative mixing). Such assortativity patterns have implications for disease spreading and control strategies. Positive assortativity can lead to more common outbreaks (lower infection rates, faster epidemic growth), but also, the disease spreads first to higher degree nodes in this network, which enables faster contact tracing and implementation of control strategies (Kiss et al., 2008). | (Newman, 2010) |

**Supplementary Table S2.** Description of network analysis terminology and parameters

| Production type classification | Classification provided by company |
|---|---|
| **Sow** | "Sow," "Sow Unit," "Isolation, Sow," "Composter, Sow," "Composter, Finishing, Sow." |
| **Nursery** | "Composter, Nursery," "Nur," "Adj," "Nursery." |
| **Wean-to-finisher** | "Wean To Finish," "Wean_to_finish," "Weantofinish," "wean-to-finisher," "Nursery; Finisher," "Finishing, Nursery." |
| **Finisher** | "Finisher," "Finishing," "Fin," "Finish," "Composter, Finishing," "Gilt Finishing," "Gf." |
| **GDU** | "Gdu," "Gilt," "Sow & Gdu." |

**Supplementary Table S3.** Number of interstate and intrastate shipments and pigs from 1st January 2020 to 15th January 2023

| Origin | Destination | Number of shipments | Number of pigs |
|---|---|---|---|
| NC | NC | 88,749 | 322,713,497 |
| IA | IA | 17,505 | 7,001,558 |
| OK | OK | 15,717 | 4,861,502 |
| TX | TX | 11,141 | 2,743,758 |
| KS | KS | 7,670 | 2,890,489 |
| IL | IL | 6,051 | 4,203,479 |
| OK | KS | 4,534 | 1,407,649 |
| PA | PA | 4,118 | 2,583,607 |
| CO | CO | 4,005 | 1,109,388 |
| IL | IA | 3,422 | 2,607,135 |
| NC | VA | 3,011 | 11,152,229 |
| NE | IA | 2,656 | 1,621,835 |
| KS | OK | 2,319 | 887,563 |
| NC | SC | 2,282 | 6,311,509 |
| NE | NE | 2,230 | 1,026,555 |
| WY | IA | 1,738 | 741,089 |
| IN | IN | 1,714 | 975,199 |
| VA | NC | 1,658 | 1,770,461 |
| OK | IL | 1,555 | 1,128,672 |
| SC | NC | 1,510 | 3,038,413 |
| OK | IA | 1,330 | 528,859 |

| | | | |
|---|---|---|---|
| OK | TX | 1,200 | 443,723 |
| TX | OK | 1,125 | 233,294 |
| TX | KS | 933 | 335,799 |
| WY | WY | 710 | 43,258 |
| IN | IL | 680 | 361,470 |
| IL | MO | 665 | 574,544 |
| VA | VA | 554 | 2,064,258 |
| CO | OK | 448 | 153,507 |
| OK | CO | 424 | 38,357 |
| CO | IA | 420 | 143,405 |
| SC | SC | 413 | 977,181 |
| NC | MO | 363 | 1,233,633 |
| NC | PA | 357 | 470,410 |
| OK | MO | 333 | 234,875 |
| NC | IA | 244 | 1,144,033 |
| CO | KS | 235 | 73,037 |
| NE | OK | 221 | 99,949 |
| OH | IN | 217 | 95,380 |
| KS | IA | 215 | 119,912 |
| IN | IA | 206 | 140,254 |
| IN | MO | 195 | 126,738 |
| MO | MO | 195 | 178,450 |
| MO | IA | 179 | 140,163 |
| NC | IN | 172 | 219,375 |

| | | | |
|---|---|---|---|
| WY | NE | 150 | 59,272 |
| IL | IN | 135 | 46,629 |
| OH | OH | 134 | 55,490 |
| KS | CO | 112 | 36,863 |
| IN | OH | 108 | 99,145 |
| PA | NY | 96 | 51,785 |
| NE | MO | 76 | 10,762 |
| PA | NC | 73 | 31,546 |
| NC | OH | 70 | 97,650 |
| IA | IL | 64 | 41,925 |
| TX | IA | 55 | 20,459 |
| PA | IN | 49 | 10,680 |
| IL | WY | 46 | 7,546 |
| IL | NE | 44 | 10,547 |
| NC | CO | 33 | 9,798 |
| TX | NC | 26 | 10,692 |
| MO | IL | 22 | 28,879 |
| NC | OK | 16 | 3,894 |
| CO | TX | 15 | 2,534 |
| IL | OK | 15 | 1,327 |
| KS | TX | 15 | 6,519 |
| IN | PA | 12 | 5,560 |
| NC | AZ | 12 | 2,156 |
| NC | UT | 10 | 2,548 |

| | | | |
|---|---|---|---|
| IA | MO | 7 | 7,485 |
| NC | IL | 3 | 1,181 |
| VA | IA | 2 | 12,628 |
| IN | WY | 1 | 274 |
| NE | IN | 1 | 119 |

**Supplementary Table S4.** Pair-wise comparison of level of com commingling between companies by farm types

| Companies | P-value | Production type |
|---|---|---|
| A - B | < 0.05 | Sow* |
| A - C | < 0.05 | Sow* |
| B - C | < 0.05 | Sow* |
| A - D | < 0.05 | Sow* |
| B - D | 0.90 | Sow |
| C - D | < 0.05 | Sow* |
| A - E | < 0.05 | Sow* |
| B - E | 0.61 | Sow |
| C - E | < 0.05 | Sow* |
| D - E | 0.75 | Sow |
| A - F | 0.07 | Sow |
| B - F | < 0.05 | Sow* |
| C - F | < 0.05 | Sow* |
| D - F | < 0.05 | Sow* |
| E - F | < 0.05 | Sow* |
| A - B | < 0.05 | Wean-to-finisher* |
| A - C | < 0.05 | Wean-to-finisher* |
| B - C | < 0.05 | Wean-to-finisher* |
| A - D | < 0.05 | Wean-to-finisher* |
| B - D | 0.90 | wean-to-finisher |
| C - D | < 0.05 | Wean-to-finisher* |

| | | |
|---|---|---|
| A - E | < 0.05 | Wean-to-finisher* |
| B - E | 0.61 | wean-to-finisher |
| C - E | < 0.05 | Wean-to-finisher* |
| D - E | 0.75 | wean-to-finisher |
| A - F | 0.07 | wean-to-finisher |
| B - F | < 0.05 | Wean-to-finisher* |
| C - F | < 0.05 | Wean-to-finisher* |
| D - F | < 0.05 | Wean-to-finisher* |
| E - F | < 0.05 | Wean-to-finisher* |
| A - B | < 0.05 | Nursery* |
| A - C | < 0.05 | Nursery* |
| B - C | < 0.05 | Nursery* |
| A - D | < 0.05 | Nursery* |
| B - D | 0.90 | Nursery |
| C - D | < 0.05 | Nursery* |
| A - E | < 0.05 | Nursery* |
| B - E | 0.61 | Nursery |
| C - E | < 0.05 | Nursery* |
| D - E | 0.75 | Nursery |
| A - F | 0.07 | Nursery |
| B - F | < 0.05 | Nursery* |
| C - F | < 0.05 | Nursery* |
| D - F | < 0.05 | Nursery* |
| E - F | < 0.05 | Nursery* |

| | | |
|---|---|---|
| A - B | < 0.05 | Finisher* |
| A - C | < 0.05 | Finisher* |
| B - C | < 0.05 | Finisher* |
| A - D | < 0.05 | Finisher* |
| B - D | 0.90 | Finisher |
| C - D | < 0.05 | Finisher* |
| A - E | < 0.05 | Finisher* |
| B - E | 0.61 | Finisher |
| C - E | < 0.05 | Finisher* |
| D - E | 0.75 | Finisher |
| A - F | 0.07 | Finisher |
| B - F | < 0.05 | Finisher* |
| C - F | < 0.05 | Finisher* |
| D - F | < 0.05 | Finisher* |
| E - F | < 0.05 | Finisher* |

* p-value <0.05 of Kruskal-Wallis Rank Sum Test.

**Supplementary Table S5.** Summary of in-loyalty and out-loyalty values by production type

| Direction | Farm type | Quantile 25% | Median | Quantile 75% | Maximum |
|---|---|---|---|---|---|
| In-going | Finisher | 0.63 | 0.71 | 0.73 | 0.94 |
| In-going | Nursery | 0.680 | 0.77 | 0.82 | 0.96 |
| In-going | Sow | 0.61 | 0.68 | 0.73 | 0.88 |
| In-going | Wean-to-finisher | 0.39 | 0.47 | 0.68 | 0.74 |
| Out-going | Finisher | 0.41 | 0.47 | 0.51 | 0.81 |
| Out-going | Nursery | 0.27 | 0.51 | 0.86 | 0.91 |
| Out-going | Sow | 0.76 | 0.81 | 0.85 | 0.91 |
| Out-going | Wean-to-finisher | 0.14 | 0.21 | 0.52 | 0.59 |
